# MERLIN-DEFICIENT iPSCs AS AN *IN VITRO* MODEL SYSTEM FOR STUDIYNG *NF2* PATHOGENESIS

**DOI:** 10.1101/2022.12.14.520389

**Authors:** Núria Catasús, Miguel Torres-Martin, Inma Rosas, Bernd Kuebler, Gemma Casals-Sendra, Helena Mazuelas, Alex Negro, Francesc Roca-Ribas, Emilio Amilibia, Begoña Aran, Anna Veiga, Ángel Raya, Bernat Gel, Ignacio Blanco, Eduard Serra, Meritxell Carrió, Elisabeth Castellanos

**Author notes:** Corresponding authors: Meritxell Carrió and Elisabeth Castellanos ( /). Equally contributed (Meritxell Carrió, Elisabeth Castellanos).

## Abstract

NF2-related schwannomatosis is an autosomal dominant syndrome that predisposes to the development of benign tumors of the nervous system. Schwannomas, particularly bilateral vestibular schwannomas (VS), are the most characteristic features of the disease. These tumors are caused by the bi-allelic inactivation of the *NF2* gene in a cell of the Schwann cell lineage. Our current understanding of the molecular pathogenesis of the *NF2* gene, as well as the development of new effective therapies is hampered by the absence of human non-perishable cell-based bearing distinct *NF2* pathogenic variants. With this aim, we generated and characterized three isogenic paired induced pluripotent stem cell (iPSC) lines with single or bi-allelic inactivation of *NF2* by combining the direct reprogramming of VS cells with the use of CRISPR/Cas9 editing. Our results show a critical function of *NF2* for the maintenance of a stable pluripotent state. However, we were able to nudge them towards the Neural Crest-Schwann Cell (NC-SC) axis by applying a 3D Schwann cell differentiation protocol. *NF2*(+/−) and *NF2*(−/−) spheroids homogeneously expressed classical markers of the NC-SC lineage. In addition, *NF2*(−/−) SC-like spheroids showed dysregulation of multiple signaling pathways already described for merlin-deficient SC, and altered in human schwannomas. Therefore, *NF2*(+/−) and *NF2*(−/−) SC-like spheroids can represent a bona fide human *in vitro* cellular model to study the role of *NF2* pathogenesis.

## INTRODUCTION

NF2-related schwannomatosis (NF2-related SWN; OMIM 101000) is a dominantly inherited autosomal syndrome characterized by the development of multiple tumors of the nervous system (1,2). The incidence of the disease is estimated to be 1 in 28,000 live births with complete penetrance by the age of 60, with 50-60% of cases due to *de novo* mutations (3). The pathognomonic feature of the disease is the development of bilateral vestibular schwannomas (VS), well-circumscribed tumors that arise from Schwann cells (SC) in the vestibulocochlear nerve, due to the bi-allelic inactivation of *NF2*. VS are responsible for hearing loss, deafness and tinnitus(4). Affected individuals may also develop multiple schwannomas in other cranial, spinal and peripheral nerves as well as intracranial and intraspinal meningiomas, and spinal ependymomas (5–8). Cutaneous and ocular manifestations are also common (9,10). Due to their number and anatomical location, these tumors, although histologically benign, are responsible for high morbidity and reduced life expectancy (11). The gold standard treatment for NF2 patients includes surveillance, surgery, radiotherapy and bevacizumab for VS (4), although this last is associated with toxicity if used for extended periods (12), so there is an important need to develop new treatments.

NF2-related SWN is caused by pathogenic variants in the *NF2* tumor suppressor gene (22q12.2), which encodes for merlin (moesin-ezrin-radixin-like) protein, a scaffold protein that interacts with transmembrane receptors and intracellular effectors to mediate mitogenic, survival, proliferation and cell-cell contact signaling pathways (13–15). Changes in merlin result in the inhibition of cell proliferation and the dysregulation of a wide variety of signaling cascades from the cell surface to the nucleus. One of the most studied roles of merlin is its involvement in the Hippo signaling pathway, by repressing YAP/TAZ nuclear translocation. Once in the nucleus, YAP and TAZ associate with the DNA-binding TEAD proteins (TEADs 1-4) to activate the expression of regulators of the cell cycle and apoptosis (16). Merlin also controls cell proliferation by contact inhibition through mechanisms that include the formation of stable adherens junctions, and the regulation of specific growth factor receptors, the focal adhesion kinase (FAK) and phosphoinositide 3-kinase (PI3K)/AKT/mTOR pathways (16). In addition, merlin is known to be involved in other signaling pathways such as the Ras/Raf/MAPK, TP53 and the Rac1-Pak1 pathways (17,18).

Studying the pathogenesis of the *NF2* gene and developing potential therapies remains challenging due to the lack of durable preclinical models that recapitulate the genetics and pathophysiology of NF2-realted tumors. Initial attempts to generate *in vivo* mouse models revealed the essential role of *NF2* during embryogenesis and early development (19). In *NF2*(+/−) mutant mice, the absence of a single *NF2* allele causes the appearance of malignant epithelial tumors and sarcomas with an aggressive tendency to metastasize (20). Conditional KO murine models can generate schwannomas and VS (21–23). In addition, several patient-derived xenograft (PDX) mouse models have also been developed, with varied levels of success in recapitulating the biology of human NF2-related SWN traits (24). In addition, *in vitro*, primary SC cultures derived from VS have been set up (25,26), although the limited lifespan of primary cultures can severely constrain their use for extended experiments. Transformed and immortalized VS-derived cell lines have been developed to overcome this limitation and have been widely used as platforms for drug screening (HEI-93, JEI-001) (27,28), although these models undergo physiological changes due to the immortalization process, severely reducing the value of any comparison with primary cells.

Induced pluripotent stem cell (iPSC) (29) models have been used in a wide range of diseases and represent an unlimited source of cells with multiple applications such as disease modelling, drug development, regenerative medicine, or gene regulation (30–33). The first NF2-related SWN model reported was an iPSC line harboring a homozygous variant in the *NF2* gene (UMi031-A-2), obtained by CRISPR/Cas9 gene-editing of an unaffected pluripotent cell line (34); and, more recently, an iPSC clone was derived from mononuclear bone marrow stem cells from a mosaic NF2-related SWN patient (35).

One of the main defining characteristics of iPSCs is their ability to differentiate into the three germ layers, and thus their potential to differentiate into any cell type involved in NF2-related SWN traits. Protocols have been developed to differentiate iPSC towards the SC lineage, passing through an intermediate multipotent neural crest stage (36–38). Using these protocols, it has been shown that iPSC-derived SCs show homogeneous expression of SC markers and the ability to myelinate axons when co-cultured with dorsal root ganglia neurons (38). Therefore, the use of iPSCs appears to be a suitable human cellular model to study the molecular pathogenesis of the *NF2* gene, and the differentiation capacity and growth behavior of the *NF2* deficient SC-derived cells that originate NF2-related schwannomas.

In the present study, we generated three isogenic iPSC lines harboring single or bi-allelic loss-of-function variants in the *NF2* gene by direct reprogramming of VS cells and the use of CRISPR/Cas9 editing. For the first time, these cells were characterized and differentiated towards the NC-SC lineage, resulting in *NF2*(+/−) and *NF2*(−/−) SC-like spheroids co-expressing classic markers of the NC-SC axis. In addition, *NF2*(−/−) SC-like spheroids showed dysregulation of several signaling pathways previously described in *NF2*(−/−) cells, providing a bona fide *in vitro* cellular model to study the role of *NF2* pathogenesis.

## MATERIALS AND METHODS

All procedures performed were in accordance with the ethical standards of the IGTP Human Research Ethics Committee (CEIC) (PI-17-250), which approved this study, and with the 1964 Helsinki declaration and its later amendments. The “ *Comisión de Garantías para la Donación y Utilización de Células y Tejidos Humanos, ISCIII*”) authorized the Project.

### Patients and Vestibular Schwannoma samples

tumor samples were obtained from NF2-related SWN patients clinically managed at the Spanish Reference Centre (CSUR) for phacomatoses HUGTiP-ICO-IGTP, diagnosed according to standard diagnostic criteria (**Supplemental Table S1**) (39). Written informed consent was obtained for iPSC generation and genomic analysis. Tumor samples were processed as described in Castellanos et al., 2018 (40).

### Reprogramming of Vestibular Schwannomas

between 0.2·10^6^-1·10^6^ of VS derived cells were reprogrammed using the CytoTune™-iPS 2.0 Sendai Reprogramming Kit (Thermo Fisher Scientific), a non-integrative cell reprogramming method holding the four Yamanaka factors (OCT4, SOX2, KLF4, and cMYC), according to manufacturer’s instructions.

### Tumor and iPSC molecular characterization

genetic analysis was performed using the customized I2HCP panel as previously described (41) or Whole Exome Sequencing (WES). The latter was performed using KAPA HyperCap technology with KAPA HyperExome Probes (Roche) according to manufacturer’s instructions and sequenced in a NextSeq instrument (Illumina). Analysis of small nucleotide variants was performed with Mutect2 and annotated with Funcotator (42). Potential SNV and small indels were detected among the original cell lines and the paired CRISPR lines, GATK 4.2.6.1 was used for the entire process (42) (**Supplemental Material and Methods**). To detect *NF2* splicing variants, tumors were studied both at RNA and DNA level. *NF2* variants were confirmed by Sanger Sequencing. SNP-array analysis was assessed using Illumina HumanOmniExpress v1 BeadChips (730,525 SNPs) according to manufacturers’ instructions and as described in Supplemental Material and Methods.

### CRISPR/Cas9 gene edition in iPSC lines

CRISPR/Cas9 editing was performed with the ArciTect ribonucleoprotein (RNP) system (STEMCELL Technologies). The designed sgRNA targeted exon 2 of the *NF2* gene (GTACACAATCAAGGACACAG) using the Synthego - CRISPR Design Tools (https://www.synthego.com/products/bioinformatics/crispr-design-tool). TransIT-X2® Dynamic Delivery System (Mirus) was used for transfection according to manufacturers’ instructions.

### iPSC culture

iPSCs were grown on growth factor-reduced Matrigel (BD Biosciences)-coated 6-well plates and cultured in mTESR Plus medium (STEMCELL Technologies). iPSCs were split using Accutase (Merk) and cells were seeded with Rock Inhibitor (STEMCELL Technologies) (1:1000) for 24h. When required, visibly differentiated cells were removed manually from the culture of *NF2*(−/−) cells. In this study, the *NF2*(+/+) FiPS line (FiPS Ctrl 1-SV4F-7) was used for all experiments as a control.

### Neural Crest (NC) differentiation

differentiation of iPSC lines into NC was performed as previously described in Menendez et al. (2013) (37) with minor modifications (38). 9×10^4^ iPSCs were seeded on matrigel-coated dishes in mTESR Plus medium. The following day, the medium was replaced with Neural Crest Differentiation Media (**Supplemental Material and Methods**) and was replaced every day. NC were maintained in this medium and split with Accutase when necessary.

### Differentiation into SC in 2D

to establish SC differentiation, 0.4×10^6^ NC cells/well were plated onto 0.1mg/mL poly-L-lysine (Sigma) and 4µg/mL laminin (Gibco) 6-well plates and cultured in SC differentiation media (SCDM) (**Supplemental Material and Methods**) as previously described (38).

### Differentiation towards SC in 3D

NC cells were detached with Accutase and, 2.25·10^6^ cells/well were seeded onto AggreWell TM800 24-well plates (Stem Cell Technologies) in 2mL SCDM (described above). The medium was changed twice a week removing 1 mL and replacing 1 mL of fresh SCDM. On days 7, 14 and 30 spheroids were collected and processed for subsequent analysis.

### Flow cytometry

accutase-dissociated cells were resuspended in PBS-0.1% BSA and incubated with p75 primary antibody and subsequently incubated with Alexa Fluor 568 secondary antibody, followed by the incubation of Hnk1 primary antibody that was detected with Alexa Fluor 488-conjugated secondary antibody. Antibodies were incubated 30 min on ice. Flow cytometry analysis was performed using BD LSR Fortessa SORP and BD FACS Diva 6.2 software.

### Immunocytochemistry

cells or spheroids were fixed in 4% paraformaldehyde in PBS for 15min at room temperature (RT), permeabilized with 0.1%Triton-X 100 in PBS for 10 min at RT, blocked in 10% FBS in PBS for 15 min at RT, and incubated with the indicated antibodies (**Supplemental Table M1**) overnight at 4°C. Secondary antibodies Alexa Fluor 488 and Alexa Fluor 568 were incubated for 1h at RT. Nuclei were stained with DAPI and images captured using LEICA DMIL6000 and LASAF software.

### RNA processing, sequencing and analysis

Total RNA extraction from iPSCs, NC cells and SC-differentiating spheroids was extracted with the 16 LEV simplyRNA Purification kit (Promega), following manufacturer’s instructions. RNA was quantified with a Nanodrop 1000 spectrophotometer (Thermo Scientific). The polyA RNA libraries were sequenced on an Illumina Novaseq 6000 in 150 bp pair-end mode. For RNAseq processing, genome GRCh37.p13 and gene annotation gencode version 19 were used. Reads were aligned with STAR version 2.7.10a (43) and count files were produced with QoRTs version 1.3.6 (44). These raw counts were used for DESeq2 differential expression (DE) analysis with Wald significance test and Benjamini & Hochberg correction (FDR) correction. A corrected p-value of 0.01 and an absolute Log2-fold change of 1 was used for considering a gene as DE. Volcano plots were performed using EnhancedVolcano (with the https://github.com/kevinblighe/EnhancedVolcano) outputs from Deseq2. To account for more direct measurement, barplots in Figure 2 were instead normalized by Trimmed Mean of M-values from EdgeR (45) and then RPKMs were calculated for every gene. The *vst* normalization from DESeq2 (46) was used for unsupervised cluster heatmaps with Euclidean metric and Ward method. PCA features were chosen using the top 2000 genes with the highest SD over vst normalization. Pathway enrichment analyses for DE genes were performed using GSEA pre-ranked test (47) using the “stat” value from Deseq2. A FDR <0.05 was considered as significant. Single sample GSEA (ssGSEA) with MSigDB version 2022.1.Hs (48) was used when comparing more than two groups of samples. Enrichr (49,50) was used only with DE genes as input. The 41 raw fastq files schwannomas for comparison were retrieved from the EGA repository with accession code EGAD00001002722, and were processed with the same methodology described above. For re-analysis of this cohort together with our set, vst values were recalculated for all of them without correction for batch.

## RESULTS

### 1. Generation of IPSC harboring single or bi-allelic pathogenic variants in the *NF2* gene

Three VS from three independent NF2-related SWN patients that met standard diagnostic criteria (39) were used for reprogramming (VS-25, VS-245 and VS-267) with the aim of obtaining iPSC clones harboring pathogenic variants present in NF2 patients. Molecular characterization of these tumors through the I2HCP panel (41) and SNP-array analysis uncovered the *NF2* alterations reported in **Supplemental Table S1** and **Supplemental Figure S1**. Except for the alteration in the *NF2* locus, no pathogenic variants were identified in any known oncogene or tumor suppressor gene. The non-integrating Sendai virus vector (SeV) encoding the 4 Yamanaka factors (29) was used to induce pluripotency of VS-dissociated cells and we determined the *NF2* genotype of the generated single-cell clones through Sanger sequencing (**Figure 1A, Supplemental Table S2**).

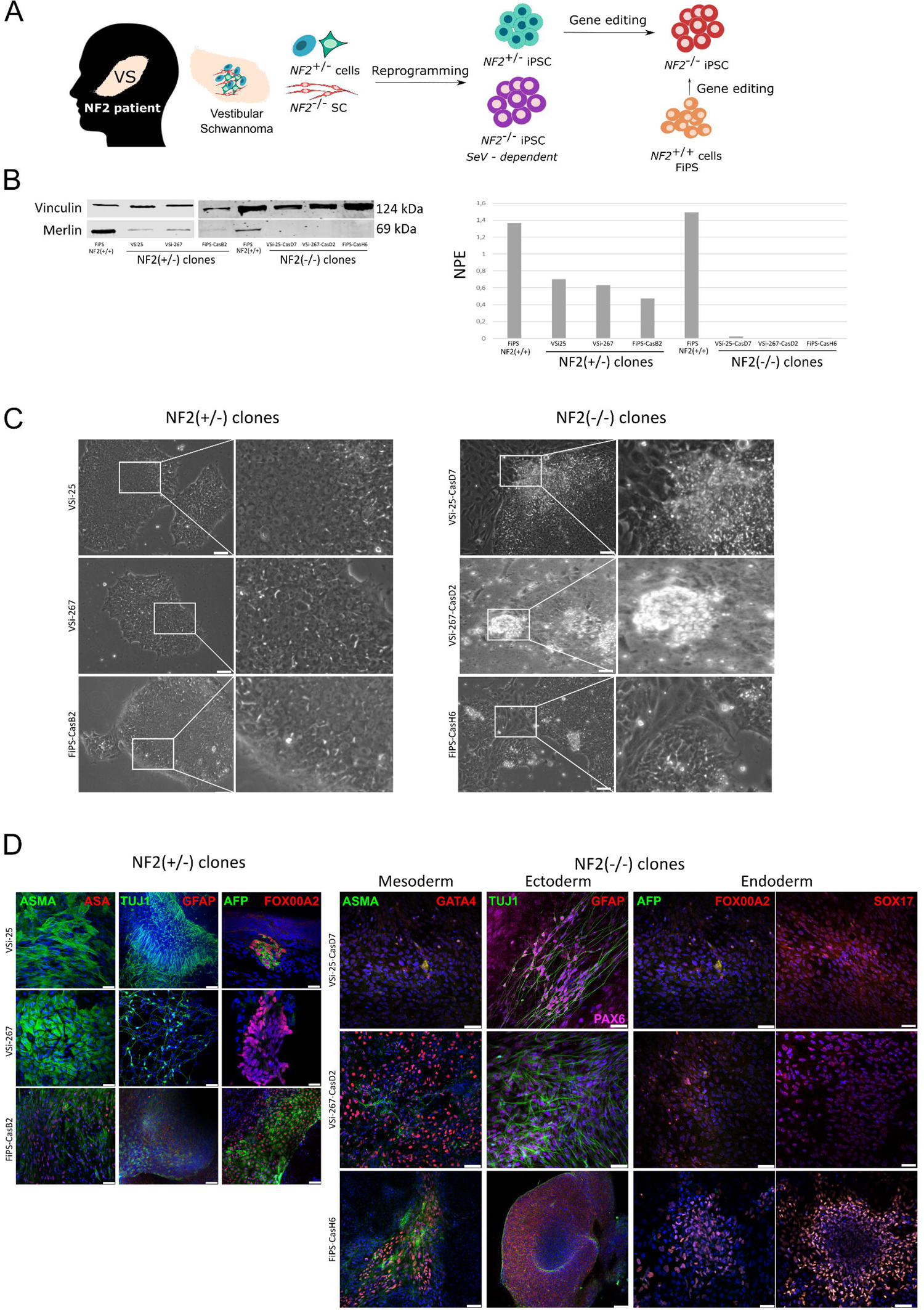

From VS-25 and VS-267, only clones carrying the germline *NF2* pathogenic variant (*NF2*(+/−)) were obtained. One clone derived from each tumor was selected for further characterization. From VS-245, a VS coming from a mosaic patient, we obtained both *NF2*(−/−) and *NF2*(+/+) iPSC clones, named VSi-245. As VSi-245(+/+) clones were reprogrammed from an unaffected cell, these were not further characterized. In contrast, all nine VSi-245 *NF2*(−/−) iPSC clones analysed sustained the exogenous expression of Yamanaka factors by the Sendai (SeV) construct for more than 40 passages after reprogramming and, therefore, were also discarded for the rest of the study (**Supplemental Figure S2**).

With the aim of achieving iPSCs with complete *NF2* inactivation, and given the low efficiencies obtained in the generation of *NF2*(−/−) clones by VS reprogramming, we used the CRISPR/Cas9 technology to edit the *NF2* gene in both, the *NF2*(+/−) VSi-25 and VSi-267 iPSCs and a control *NF2*(+/+) iPSC line (FiPS Ctrl 1-SV4F-7, registered in the Spanish National Stem Cell Bank/ ESiO44C in https://hpscreg.eu/)). Thus, using CRISPR/Cas9 RNP complexes, we targeted exon 2 of the *NF2* gene and obtained single-cell *NF2*(−/−) iPSC clones from VS-derived lines (named 25-CasD7, when derived from VSi-25, and 267-CasD2 when derived from VSi-267) and isogenic *NF2*(+/−) and *NF2*(−/−) iPSC clones derived from the control line (named FiPS-CasB2 and FiPS-CasH6, respectively). Pathogenic variants in *NF2* were fully characterized through PCR and Sanger sequencing and the *NF2* coding region was cloned to confirm compound heterozygous clones (**Table 1**). In addition, all *NF2* genotypes were confirmed through a merlin western blot (**Figure 1B**). From this point on, we worked on the characterization of three pairs of isogenic iPSC lines: three with one truncating variant in *NF2* and three lines carrying the bi-allelic inactivation of *NF2* (two pairs VS-reprogrammed and one from a control iPSC line (**Table 1**)).

**Table 1.**
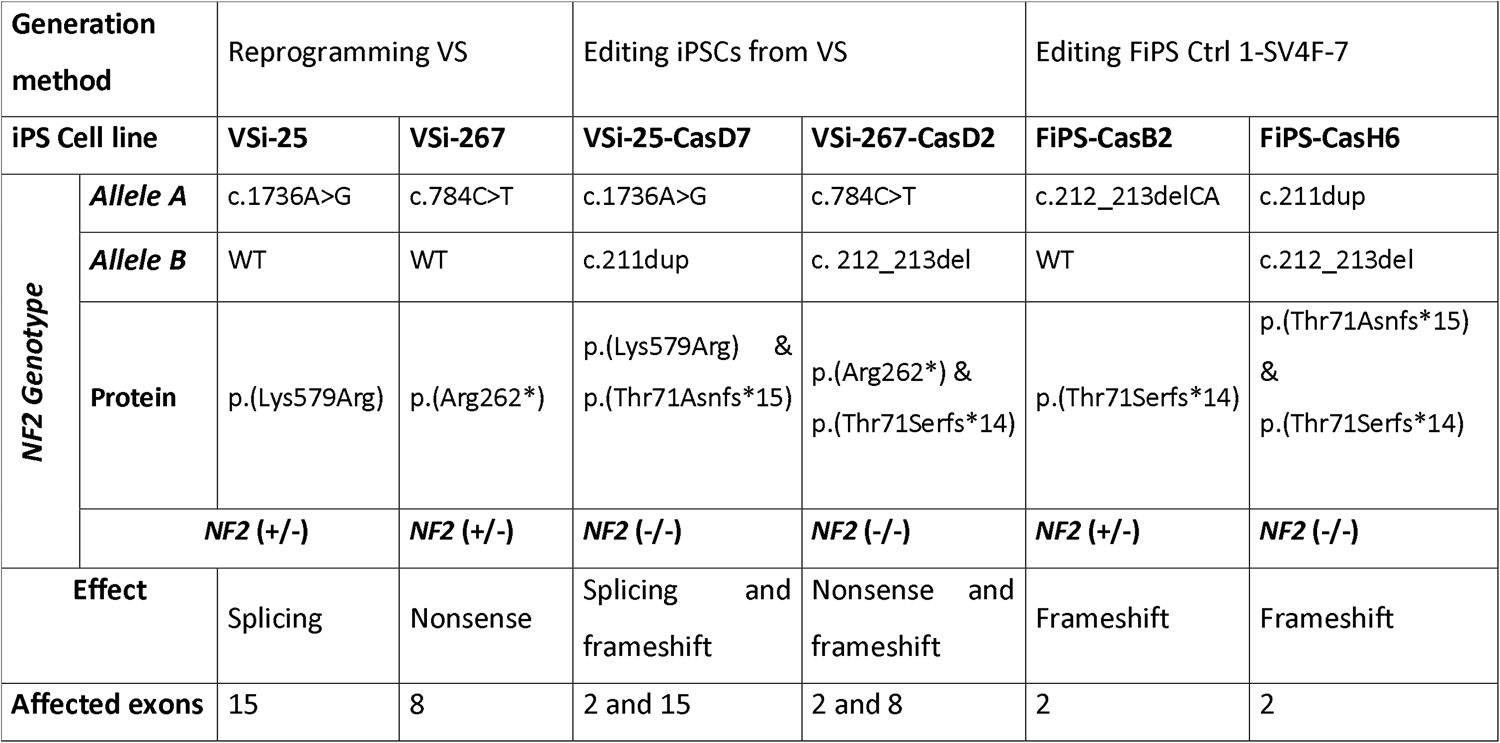
iPSCs lines information.

iPSCs: induced pluripotent stem cell; VS: vestibular schwannoma; FiPS; fibroblasts induced pluripotent stem cell; Ctrl: Control; WT: wild-type

#### 1.1. Characterization of *NF2* (+/−) and *NF2* (−/−) iPSC

All clones analysed expressed cell surface proteins and transcription factors associated with pluripotency: Oct4, SSEA-3, SSEA-4, Nanog, Sox2, Tra-1-60 and Tra-1-81; were positive for alkaline phosphatase staining; and showed karyotype stability after at least 20 passages (46, XY) (**Supplemental Figure S3, A-C**). Although all iPSC clones expressed the typical pluripotent markers, we could observe differences in clone morphology between *NF2*(+/−) and *NF2*(−/−) cell lines. Whereas *NF2*(+/−) showed classical iPSC colony morphology, edited *NF2*(−/−) iPSCs presented less compact colony formation, eventually aggregating in the center of the colony, growing upwards and undergoing major spontaneous differentiation compared with the iPSC control line (**Figure 1C**). Furthermore, *NF2*(+/−) iPSCs showed the capacity to differentiate into the three primary germ layers *in vitro* (mesoderm, endoderm, and ectoderm) through embryoid bodies (EBs) formation. In contrast, *NF2*(−/−) lines could not be differentiated using standard EBs generation protocols and required direct differentiation to acquire expression of the three germ layers (**Figure 1D, Supplemental Table S3**).

Genomic characterization of *NF2*(+/−) VSi-25 and VSi-267 iPSCs selected clones by SNP-array analysis and WES showed no differences with respect to the tumor of origin, with the exception of the absence of the the second hit in the *NF2*(+/−) lines (**Supplemental Figure S1**). Generated *NF2*(−/−) iPSC lines revealed no pathogenic off-target alterations by WES analysis of the edited lines (**Supplemental Table S4**).

We have therefore generated three *NF2* iPSCs isogenic pair lines (*NF2*(+/−) and *NF2*(−/−)) with three different genomic backgrounds and cells of origin, providing an unlimited source of cells, consisting of a biological triplicate system. The phenotype observed in all three *NF2*(−/−) lines, which showed relevant phenotypic and differentiation abnormalities, suggests that the *NF2* gene may play a role in maintaining a purely pluripotent state.

### 2. *NF2* iPSC differentiation towards the NC-SC lineage

Given that the cells that initiate schwannoma formation in NF2-related SWN are *NF2*(−/−) cells derived from the SC lineage, we applied a differentiation protocol towards the NC-SC axis. As a first step, we generated NC by applying a chemically defined differentiation protocol as previously reported (37). After ten days of NC differentiation, cells achieved NC morphology (**Figure 2A**). Cell identity was assessed by flow cytometry of NC markers, NGFR (Nerve Growth Factor Receptor) (p75) and HNK1 (CD57), at early passages of the NC differentiation process (p4) and in a mature NC state (p8). *NF2(+/+)* an *NF2(+/−)* cells showed high expression of both markers already at p4 which increased up to more than 80% at a mature NC state. *NF2*(−/−) cells already showed some expression of p75 and Hnk1 at the iPSC stage which increased at p4, but did not acquire the same percentages of p75 and Hnk1-positive cells at the mature NC stage (65-80%), compared with their controls *NF2(+/+)* and *NF2(+/−)* counterparts (**Figure 2B**). Immunocytochemistry and transcriptome analysis (RNA-seq) of NC derived cells showed that all lines expressed NC markers Sox10, TFAP2A and p75 and showed no expression of pluripotency markers POU5F1 (Oct4), Nanog and Sox2 (**Figure 2C and 2D**). However, some *NF2 (−/−)* NC cells were also expressing S100B at both the RNA and protein levels, which was consistent with the heterogeneity detected by flow cytometry assays. Nonetheless, although *NF2*-deficient lines exhibited a distinct phenotype, they could be differentiated towards a NC-like identity, were able to maintain self-renewal capacity for at least 18 passages and could be cultured after several freeze-thaw cycles.

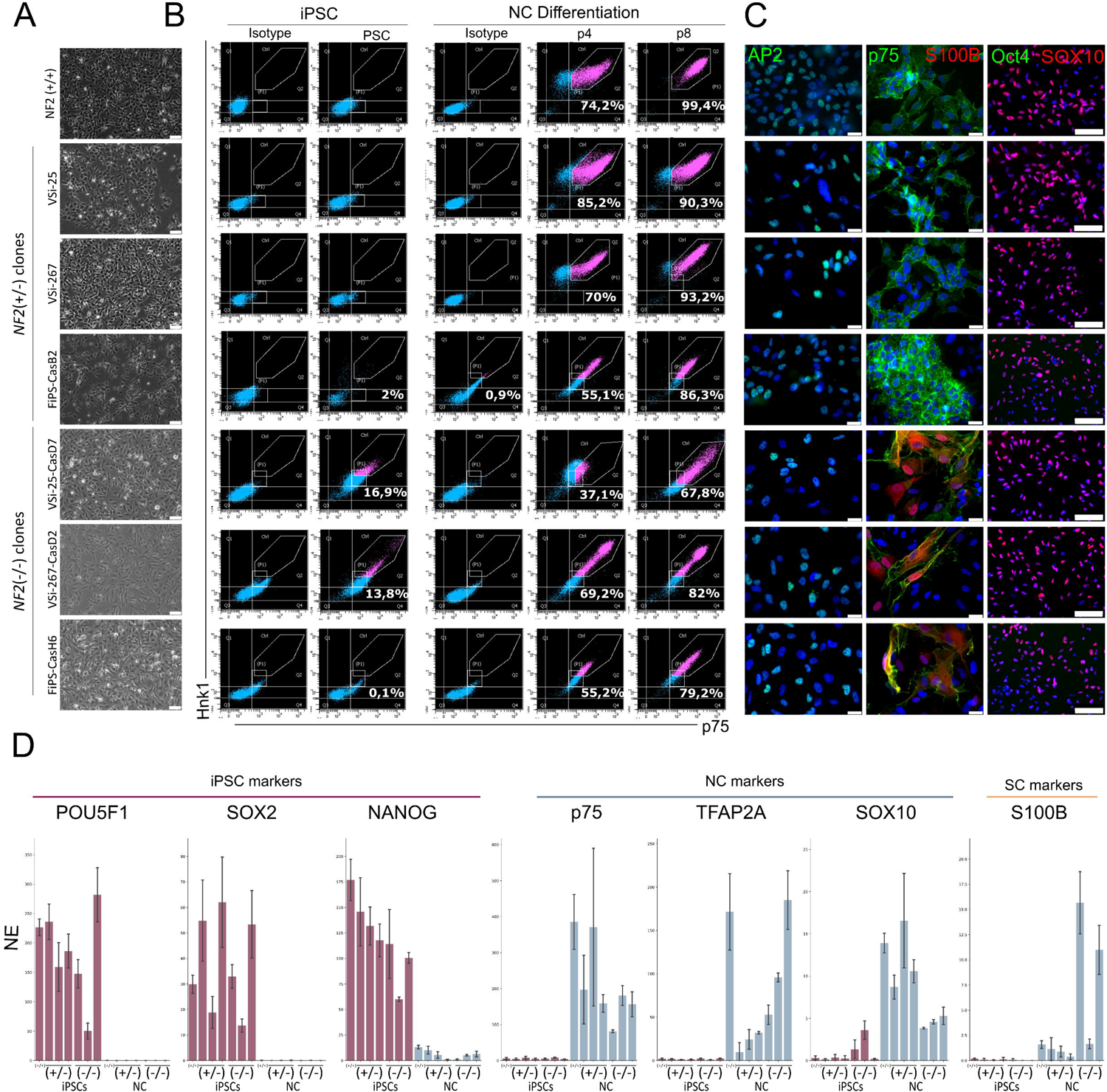

A last step in the differentiation towards SC, consisted of inducing the differentiation of the established NC cultures into a SC state. Thus, NC cells were cultured in SC differentiation medium (SCDM) for 30 days, as previously described (37,38). However, five days after inducing SC differentiation, *NF2*(+/−) and *NF2*(−/−) cells showed difficulties in maintaining adherence to culture dishes, in contrast to the control line, which after seven days of differentiation already showed a more elongated spindle morphology resembling SC-like cells (**Supplemental Figure S4**). We reasoned that the poor adherence capacity observed compromised their ability to differentiate towards a SC identity in the standard 2D stablished conditions, and therefore, we tested their capacity to differentiate in 3D conditions, following a previously established protocol (51) with minor modifications.

#### 2.1 NC-SC lineage marker analysis on *NF2*(+/−) and *NF2*(−/−) spheroids

To evaluate the SC differentiation capacity of *NF2*(+/−) and *NF2*(−/−) cell lines in 3D, we studied the transcriptome at three time points (7, 14 and 30 days) of the differentiation process. Principal component analysis (PCA) revealed that iPSCs, NCs and SCs triplicates clustered according to cell type, regardless of genotype or differentiation state (**Figure 3A**).

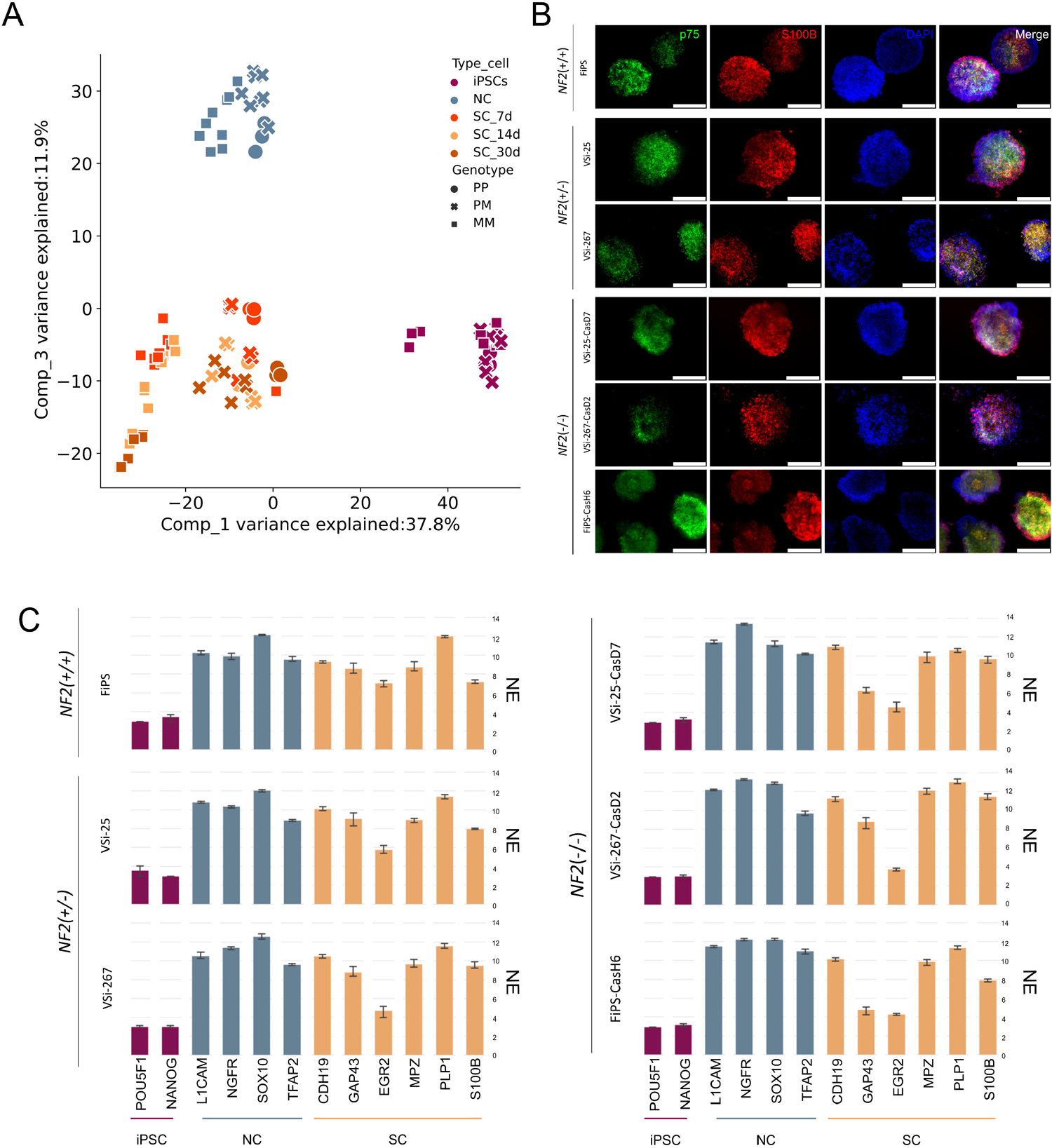

At the RNA level, we observed that *NF2*(+/+) differentiating spheroids expressed typical SC lineage markers (S100B, MPZ or PLP1) already at day 7. At day 14 of differentiation, *NF2*(+/+) spheroids were positive for p75 and S100B markers identified by immunostaining (**Figure 3B**) and expressed SC markers closely resembling those expected in SCs (**Supplemental Figure S5**). However, at day 30 of differentiation the expression of some SC markers decreased (SOX10, PLP1) whereas some central nervous glial markers appeared (FABP7, ASTN1), suggesting loss of SC commitment. For this reason, we decided to analyse samples at day 14 of differentiation (**Supplemental Figure S5**). *NF2*(+/−) VSi-25 and VSi-267 cell lines formed spheroids that stained positive for p75 and S100B markers, as did the three *NF2*(−/−) cell lines (**Figure 3B**). However, we were unable to obtain spheroids from the FiPS-CasB2 *NF2*(+/−) edited cell line despite several attempts (**Supplemental Figure S6**).

RNAseq analysis revealed that *NF2*(+/−) and *NF2*(−/−) cell lines expressed high levels of SC markers such as CDH19, GAP43, MPZ, PLP1 or S100B at 14 days of differentiation, in a similar way to the *NF2*(+/+) control line, indicating that NC cells were progressing from a precursor state into a committed SC phenotype (**Figure 3C**). Finally, we used a recently published *in vitro* NC-SC expression roadmap (50) to analyze the *NF2*(−/−) cell lines (51) and they displayed a very similar NC-SC expression roadmap to that previously determined for *NF2*(+/+) differentiating SCs in 2D (**Supplemental Figure S7**) (51). Overall, we successfully obtained *NF2*(+/−) and *NF2*(−/−) SC-like spheroids, that express NC-SC lineage markers at the RNA and protein levels, by applying 3D SC differentiation conditions in three isogenic paired lines.

### 3. Merlin-related pathway analysis in the *NF2*(−/−) SC-like cells

To better characterize *NF2*(−/−) SC-like spheroids and the effect of the absence of *NF2* on them, we performed differential expression analysis of the distinct genotypes at 14 days of SC differentiation stage. Only 125 genes were differentially expressed (DEG) when comparing *NF2*(+/+) and *NF2*(+/−) genotypes; most of them related to cellular polarity, cell adhesion and to the mTOR-pI2K-Akt signalling pathway; and some of them directly regulated by merlin, such asCHL1(52)(**Supplemental Figure S8A-B**). A higher number of DEG were found when comparing the *NF2*(+/+) and *NF2*(−/−) (**Supplemental Figure S8C**), as well as when comparing *NF2*(+/−) and *NF2*(−/−) (2874 and 3447 respectively) (**Figure 4A**), indicating that the second hit in *NF2* is the driver of the major expression changes in these cells.

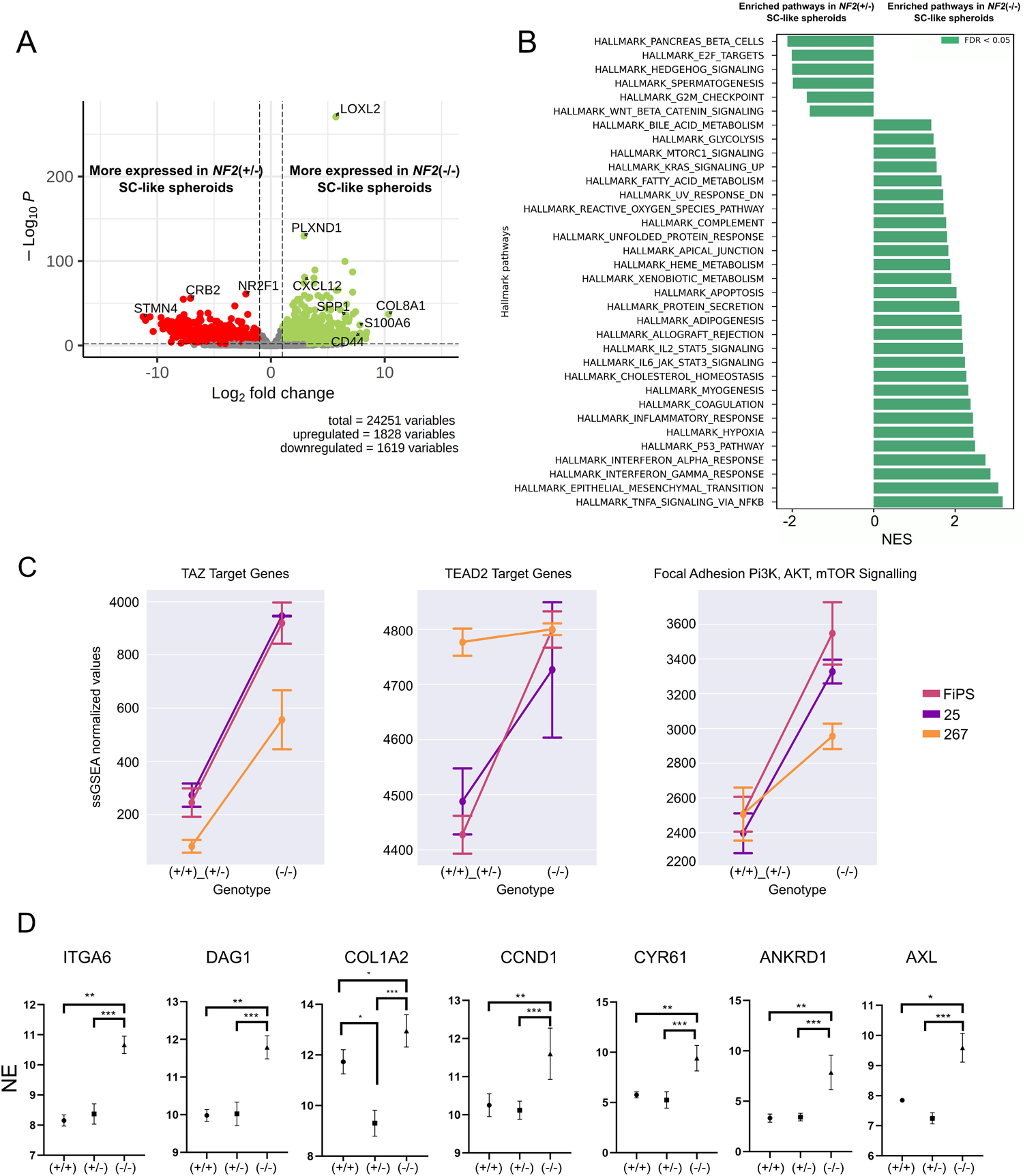

As it was not possible to obtain *NF2*(+/+) lines from SWN-related NF2 patients, we focused on comparing *NF2*(+/−) and *NF2*(−/−) SC-like spheroids and further investigated DEG by functional enrichment analyses. These showed significantly enriched Hallmark signaling pathways in *NF2*(−/−) SC-like spheroids, including mTORC1, NFKβ, p53, Hedgehog, and IL6-JAK-STAT3, among others, most of them directly or indirectly regulated by merlin. Other altered signaling pathways were Epithelial Mesenchymal Transition (EMT), cholesterol homeostasis and fatty acids metabolism, ROS signaling, hypoxia and apoptosis, Wnt/β-cantenin and finally (α and γ) interferon response (**Figure 4B**). Similar results were obtained when comparing *NF2*(+/+) vs the *NF2*(−/−) SC-like spheroids (**Supplemental Figure S8D**).

Further analyses were performed using single sample gene set enrichment analysis (ssGSEA) to test whether signaling pathways previously described in merlin-deficient cells were systematically altered in all *NF2(−/−)* SC-like spheroids. First, we examined the levels of TAZ target genes since their expression is negatively regulated by merlin. As expected from merlin deficiency, TAZ target genes were upregulated in *NF2*(−/−) SC-like spheroids. Similar results were observed for TEAD2 target genes, although we observed variability between cell lines (**Figure 4C**). Expression levels of genes regulated by the focal adhesion kinase (FAK) and phosphoinositide 3-kinase (PI3K)/AKT/mTOR pathways, activated in the absence of merlin, were upregulated in all three *NF2*(−/−) SC-like spheroid lines. (**Figure 4C**).

We also examined the expression of specific genes known to be altered in both *NF2*(−/−) primary SCs and schwannomas and compared their normalized expression levels between the three genotypes. Alterations were found in laminin receptor α6 integrin (Itga6), α-dystroglycan (Dag1), and collagen (Col1a2), consistent with an alteration of basal cytoskeletal organization in *NF2*(−/−) SC-like spheroids when comparted to *NF2*(+/+) and *NF2*(+/−) spheroids (**Figure 4D**). In addition, analysing the status of the Hippo pathway signaling, we were able to confirm that some of the major YAP target genes described in SC, involved in cell growth, proliferation and migration, were significantly up-regulated in SC-like spheroids lacking *NF2* (*CCND1, CYR61, ANKRD1, AXL*) (**Figure 4D**).

Altogether, these findings showed that the alterations and differentiation properties of *NF2*(−/−) iPSC-derived SC-like spheroids could be attributed to the lack of merlin. Moreover, these results highlight a strong correlation between the previously described altered signaling pathways and gene expression profile in merlin-deficient SC and that observed in *NF2*(−/−) iPSC-derived SC-like spheroids at 14 days of differentiation, indicating that these cells could represent a genuine *in vitro* human NF2 SC model system.

GSEA analysis comparing *NF2*(+/−) and *NF2*(−/−) SC-like cells revealed that other relevant merlin targets previously found to be altered in schwannomas were also upregulated in the *NF2*(−/−) spheroids, such as the previously mentioned MEK, PI3K-AKT/mTOR, TP53, TNFα-NFκB signaling, or *ERBB2* signaling pathways, inflammatory response as well as the EMT (**Figure 4B**). We confirmed this by specifically interrogating expression levels of *MAPK1*, *KRAS*, *AKT*, *TP53*, *NFKB* and *VEGFA* (**Supplemental Figure S9A**). These analyses indicated that *NF2*(−/−) SC-like cells could recapitulate profile signatures observed in NF2-related schwannomas.

To further explore this hypothesis, we compared the transcriptome of these cells at 14 days of differentiation with published transcriptomes of 41 schwannomas (EGAD00001002722) (53). PCA revealed that the second component was driven by the *NF2* genotype (**Supplemental Figure S9B**), thus, we extracted the top 500 genes with higher weight for each feature, and selected sub-clusters where schwannomas and *NF2*(−/−) SC-like cells showed a similar pattern of expression compared to cells harboring at least one copy of *NF2* (**Figure 5A**). EnrichR pathway analysis of these 500 genes showed 160 (out of 273) of them were upregulated in *NF2*(+/−) and *NF2*(+/+) SC-like cells and were enriched in cell motility and extracellular organization, therefore related to merlin’s function (**Figure 5B**). On the other hand, 113 genes in merlin deficient schwannomas and SC-like cells were enriched in axonogenesis and neurogenesis (**Supplemental Figure S9C**). These results suggest that schwannomas and *NF2*(−/−) SC-like spheroids share a common pattern of gene expression for key pathways related to *NF2* biology compared to wild type cells.

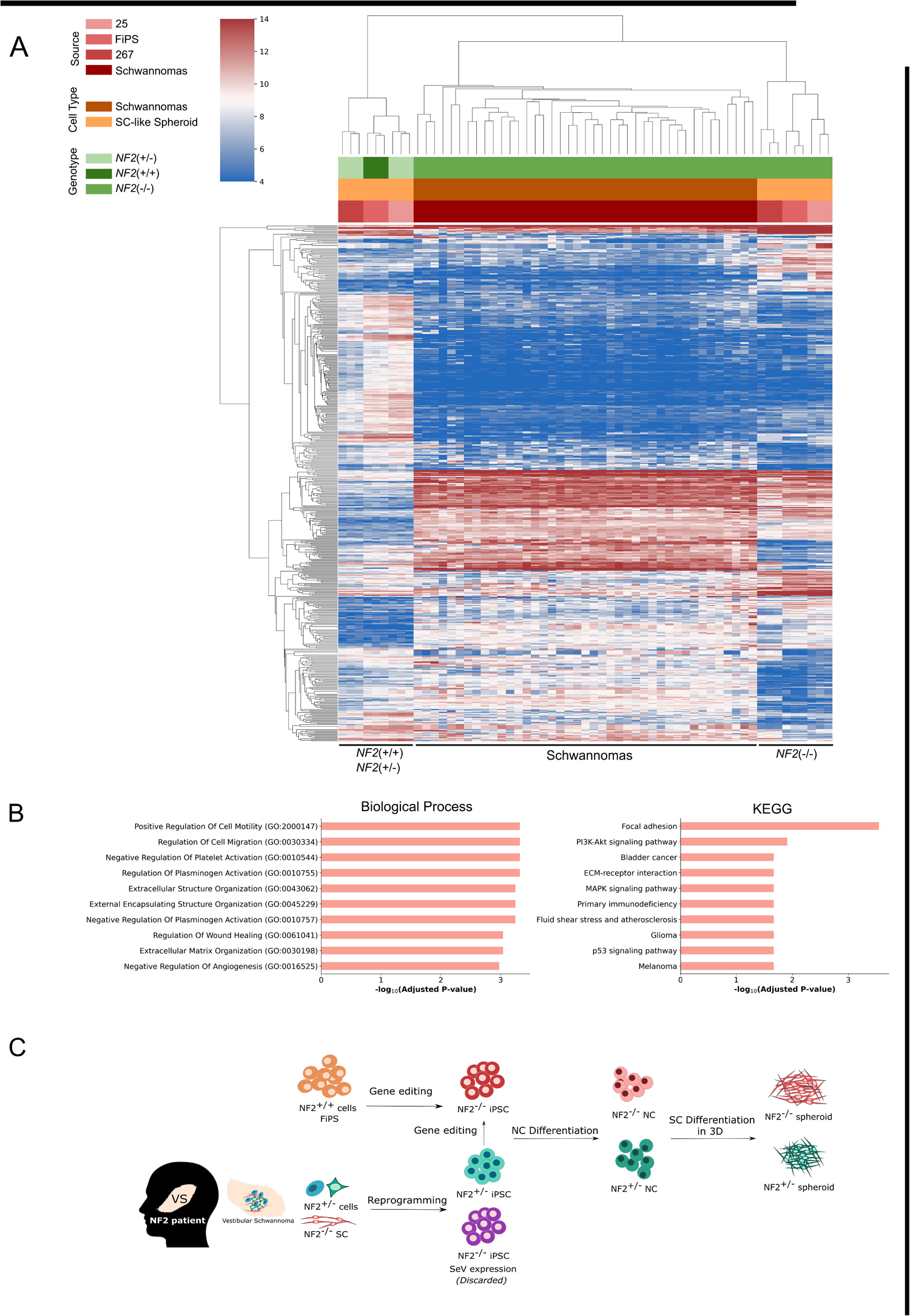

Overall, we successfully applied 3D SC differentiation conditions to *NF2*(+/−) and *NF2*(−/−) to obtain spheroids expressing NC-SC lineage markers at RNA and protein level (**Figure 5C**). In addition, gene expression of iPSC-derived merlin-deficient differentiating SCs showed that these cells constitute a bona fide cellular model for the study of the *NF2* role in Schwann cells, and potentially in any cell type associated with *NF2* pathogenesis.

## DISCUSSION

In this work we show the generation of an unperishable human iPSC-based model for the study of the molecular pathogenesis of the *NF2* gene. We have differentiated the generated isogenic iPSCs towards the NC-SC lineage and shown that *NF2*(−/−) SC-like spheroids share gene expression patterns with *NF2*(−/−) cells and schwannomas.

We first aimed to generate *NF2*(−/−) iPSCs from VS reprogramming to obtain cell lines that recapitulate the genetic background of a VS constituent cell. Our results showed low efficiencies in reprogramming NF2-deficient VS cells and we were not able to generate *NF2*(−/−) iPSCs through this process, hypothesizing that *NF2* could have a relevant role in achieving pluripotency due to its essential role in embryogenesis and early stages of development (19), which is also consistent with the reported essential function of *NF2* in haploid human iPSCs and leukemia cells (54,55). Reprogramming resulted in the generation of two *NF2*(+/−) lines (VSi-25 and VSi-267), and their *NF2* deficient isogenic cell lines (25-CasD7, 267-CasD2) were obtained by CRIPR/Cas editing. A third isogenic paired iPSC line was obtained by editing the control FiPS iPSC line (*NF2*(+/−) FiPS_CasB2 and *NF2*(−/−) FiPS-CasH6). Thus, in this study we generated a triplicate of paired *NF2*(+/−) and *NF2*(−/−) iPSC lines by combining direct reprogramming of VS cells and *NF2* gene editing, as has been recently reported (34). In our hands, the three *NF2*(−/−) iPSC lines required extensive experimental care, showed aberrant colony morphology and were susceptible to spontaneous differentiation. No differences were observed between the three lines, hinting that observed phenotypes are due to the lack of merlin and not to the iPS cell of origin. All these factors taken together point to an essential role for *NF2* in the maintenance of pluripotency of diploid cells. The three isogenic paired iPSC lines generated in this study represent a valuable and non-perishable source of cells with single or bi-allelic inactivation of *NF2*, which could be used as a NF2-cell based model to develop further assays and to study the role of *NF2* in human cells (56).

Here, we showed that both *NF2*(+/−) and *NF2*(−/−) iPSCs have the capacity to differentiate into a NC identity, although *NF2*(−/−) NC cultures exhibited an unstable NC phenotype, with some cells undergoing spontaneous differentiation and expression of S100B and requiring a higher confluence in culture to maintain their NC identity, consistent with the role of merlin in cytoskeleton maintenance and control of proliferation by cell-cell contact inhibition (57–59). Also, in line with these findings, in the process of differentiation of NCs into SCs, we observed that both *NF2*(+/−) and *NF2*(−/−) cells were easily detached from the cell culture surface, eventually leading to cell death and preventing the generation of SC-like cultures. However, and most importantly, we overcame this limitation by successfully generating *NF2*(+/−) and *NF2*(−/−) spheroids growing in suspension, as had been previously reported for other SC-based spheroids (51). These spheroids expressed classical NC-SC markers after seven days in SCDM and preserved a previously characterized *in vitro* NC-SC differentiation expression roadmap as control iPSCs (51). Therefore, for the first time, we have achieved merlin-deficient SC-like cells. *NF2*(−/−) SC-like spheroids showed deregulation of several signaling pathways as previously described in *NF2*(−/−) SC. As expected for merlin deficiency the Hippo pathway was downregulated and YAP/TAZ targets were overexpressed (*Cyr61, AXl1, ANKRD1, CCND1*) (60), while mTORC1 signaling was upregulated (61), as previously described in schwannomas, meningiomas and arachnoidal merlin-deficient cells, (60), consistent with the hypothesis that deregulation of mTORC1 activation, in addition to the Hippo pathway, underlies the aberrant growth and proliferation of NF2-deficient cells (62). Differentiated *NF2*(−/−) SC-like cells also showed upregulation of well-known merlin-related cytoskeletal organization markers, such as Itga6, Dag1 and Col1a2 (63). Furthermore, *NF2*(−/−) SC-like cells overexpressed inflammatory response, ROS and NFkB-related pathways, most probably by Rac1 deregulation as described in NF2-tumor models and merlin-deficient cells (53,64–67). Finally, merlin-deficient SC-like cells also showed upregulation of TP53, VEGF and epithelial-to-mesenchymal transition signaling, which is known to be deregulated in schwannomas (17,53,68–72). These results confirmed the strong correlation between the described and observed changes in signaling pathways and gene expression due to merlin deficiency, suggesting that the cells developed in this study could be a reliable *in vitro* human NF2 SC model system to study the molecular pathogenesis of *NF2*. In addition, because there are expression patterns described in schwannomas that reproduce in merlin-deficient SC-like spheroids; if they were validated *in vivo*, these cells would be an *in vitro* cellular model for schwannoma cells, which could be used as a platform for drug screening.

## CONCLUSION

In this study, we generated three isogenic paired *NF2*(+/−) and *NF2*(−/−) iPSC lines through reprogramming VS and *NF2* editing. According to our results, the *NF2* gene seems to play a relevant role in the maintenance of pluripotency. For the first time, iPSCs with different *NF2* genotypes were differentiated into NC-SC lineage cells, although *merlin*-deficient iPSCs exhibited an altered differentiation capacity. The application of a 3D SC differentiation protocol resulted in the successful generation of *NF2*(+/−) and *NF2*(−/−) SC-like spheroids expressing NC-SC lineage markers. In addition, *NF2*(−/−) SC-like spheroids showed dysregulation of several signaling pathways previously described in merlin-deficient Schwann cells, such as mTOR and Hippo-YAP, amongst others. Therefore, *NF2*(−/−) SC-like spheroids constitute a valuable resource for studying *NF2* biology in Schwann cells and potentially in any cell type associated with *NF2* pathogenesis.

## ACKNOWLEDGEMENTS

We thank the HGTP Clinical Services and staff for their collaboration in generating and collecting patient samples. We thank the IGTP Flow Cytometry and IGTP Genomics core facilities and their staff for their contribution and technical support. We would like to acknowledge the constant support of the different NF lay associations: *Asociación de Afectados de Neurofibromatosis*, Chromo22 and ACNefi. This study has been funded by the Chromo22 Patients association; the *Instituto de Salud Carlos III* through the project PI20/00215 (Co-funded by European Regional Development Fund “A way to make Europe”), and through the project AC22/00033, partner of the EJP RD. The EJP RD initiative has received funding from the European Union’s Horizon 2020 research and innovation program under grant agreement N°825575”; funded also by *Fundació La Marató de TV3* (126/C/2020), the Children’s Tumor Foundation (CTF-2019-05-005, CTF-2022-05-005), Fundación Proyecto Neurofibromatosis, the Catalan NF Association (AcNeFi), and the Government of Catalonia (SGR-Cat 2021 - 00967).

## AUTHOR CONTRIBUTIONS

**NC**: Collection and/or assembly of data; Data analysis and interpretation; Manuscript writing; **MT-M, BG:** Data analysis and interpretation; **AN, BK, IR, GC-S, BA:** Collection and/or assembly of data; **HM, AV, AR, IB:** Scientific input; **FR-R, EA** : Provision of study material or patients; **ES, MC, EC:** Conception and design; Financial support; Manuscript writing; Final approval of manuscript.

## DATA AVAILABILITY STATEMENT

The authors confirm that the data supporting the findings of this study are available within the article [and/or] its supplementary materials. Raw data for WES and RNAseq is allocated at the European Geno-Phenome Archive (EGA) with accession number (XXXXXXXXX) and will be made available to external researchers upon reasonable requests.

