## Supplementary Material and Figures for "MERLIN-DEFICIENT iPSCs AS AN *IN VITRO* MODEL SYSTEM FOR STUDIYNG *NF2* PATHOGENESIS"

**Supplemental Material and Methods**

**Tumor collection and cryopreservation:** tumor samples were placed in DMEM medium (Gibco) with 1x Glx (Gibco) and 1x normocin antibiotic cocktail (InvivoGene) after surgery resection and cryopreserved in 10% DMSO (Sigma) + 90% FBS until processed.

**Tumor processing:** VS were digested with 160U/mL Collagenase type 1 and 0.8U/mL Neutral protease (Worthington, Lakewood, NJ) for 16 h at 37^o^C. Dissociated cells were seeded on 0.1 mg/mL Poly-L-lysine (Sigma) and 4 µg/mL laminin (Gibco)-coated dishes in Schwann cell medium (SCM) and maintained at 37^o^C and 10% CO_2_ atmosphere. SCM consists of: DMEM (Gibco) with 10% FBS, 500 U/mL penicillin/500mg/mL streptomycin (Gibco), 0.5mM 3-iso-butyl-1-methilxantine (Sigma), 2.5 µg/mL insulin (Sigma), 10nM heregulin-b1 (PeproTech), and 0.5µM forskolin (Sigma).

**SNP-array analysis**: Raw data was processed with Illumina Genome Studio v2011.1 with the Genotyping module v1.9.4. All samples were analysed independently and treated as unpaired samples.

**Variant analysis:** Human Genome Variation Society (www.hgvs.org) nomenclature guidelines were used to name the mutation at the DNA level, its effect at the mRNA level, and the predicted resulting protein. The first nucleotide of the ATG translation initiation codon is denoted position þ1 according to the *NF2* mRNA sequence NM_000268.3 5. and NM_016418.

**CRISPR/Cas9 gene edition in iPSC lines analysis:** gDNA from single-cell clones was analysed to screen *NF2* mutations. In order to study cis o trans variants in *NF2, NF2* cDNA was cloned using Gateway Technology (Invitrogen). iPSC lines obtained from reprogrammed vestibular Schwannomas (VS-25, VS-245 and VS-267) and the control *NF2*(+/+) line used in this study (FiPS Ctrl 1-SV4F-7) obtained from the Spanish National Stem Cell bank (BNLC) were used to obtain isogenic *NF2*(-/-) iPSC lines.

- **Methods for SNV and indel off target analysis:** Materials including genome, indel and SNP databases were retrieved from ftp.broadinstitute.org at /bundle/b37. For alignment, bwa 0.7.17 (1) was used under hg19 genome assembly. Following GATK Best Practices (2), aligned bam files were marked for duplicated reads and Base Quality Score Recalibration was applied to each specimen. Afterwards, Mutect2 with orientation bias correction was used for SNV and indel calling for each pair pre/post CRISPR cell line call. Variants with TLOD > 20, counts > 30 and AF > 30% were considered as a potential CRISPR off-target candidates. Conflictive and suspicious calls were manually reviewed using IGV (3). For potential CRISPR off-target sites created from our Cas sequence, we used cas-offinder (4) online version (<http://www.rgenome.net/cas-offinder/>) with lenient thresholds (max DNA/RNA Bulge Size 2 and max mismatch of 3). These sites were flanked +/-1000bp for subsequent analysis. Pybedtools (5) the Python wrapper for Bedtools (6) was used to intersect the detected SNP/indels sites with the potential off target sites. We consider at least 1bp of overlap as an off-target. In order to annotate the variants, GATK’s Funcotator with source version 1.7.20200521s was used.

**iPSC characterization**

**Immunohistochemistry of pluripotency-associated markers:** iPSCs were fixed with 4% paraformaldehyde (PFA), blocked and permeabilized with TBS + 0.5% Triton X-100 + 6% donkey serum. Primary antibodies were incubated overnight in TBS + 0.1% Triton X-100 + 6% donkey serum. Secondary antibodies were incubated for 2h at 37ºC. Nuclei were stained with 4',6-diamino-2-fenilindol (DAPI). Antibodies are listed in Supplemental Table M1. Confocal images were taken using Leica TSC SPE/SP5 microscopes.

**Karyotype determination:** Karyotype of iPSCs was evaluated by G banded metaphase karyotype analysis, at the Hospital San Joan de Deu (Barcelona) and at the Cytogenetics Platform of the Haematology Department at Germans Trias I Pujol Hospital (Badalona).

**Alkaline Phosphatase activity**: Alkaline Phosphatase Blue Substrate Solution (Sigma) was used to demonstrate iPSC alkaline phosphatase activity. iPSC karyotype was assessed by treating cells with colcemid (Gibco) and processed as described (7).

RT-PCRs were performed to confirm the absence of the Sendai reprogramming virus and transgene expression (Supplemental Table M2).

**Merlin Western Blot:** Cells were lysed with RIPA buffer (50 mM Tris-HCl (pH 7.4), 150 mM NaCl, 1mM EDTA, 0.5% Igepal CA-630) supplemented with 3mM DTT (Roche), 1mM PMSF (Fluka), 1mM sodium orthovanadate (Sigma), 5mM NaF (Honeywell), 10 ug/ml leupeptin (Sigma), 5ug/ml aprotinin (Sigma) and 1xPhosSTOP (Roche). 50 µg of protein extracted from iPSC cell lines was loaded to SDS-PAGE (150V) and transferred onto PVDF membranes (1 hour 350 mA at 4^o^C). Odyssey Blocking Buffer TBS (LI-COR) was used to block the membranes. Primary antibodies were incubated at 4^o^C overnight. Membranes were later incubated with IRDye 680LT and IRDye 800CW secondary antibodies (1:1000 dilution, LI-COR) for 1h at room temperature and scanned and analysed using the Odyssey Infrared Imaging System (LI-COR). Western Blot primary antibodies used were α-NF2: NF2/Merlin antibody (ab88957) α-mouse (Abcam) to target merlin and α-vinculin [EPR8185] (ab129002) – α – rabbit (Abcam) was used to normalize protein expression between samples.

**iPSC differentiation into the three germinal layers through embryoid body (EB) formation:** iPSC colonies were lifted as usual with EDTA and transfer in a 96 well plate using a multichannel pipette in mTeSR-1 medium (Stem Cell Technologies). The 96 wells plate were centrifuged at 800g for 10 min and incubated at 37ºC and 5% CO2 for 24h. Then, early EBs were transferred to an ultra-low attachment plate in mTeSR-1 for additional 24h. After this time, EBs were transferred to matrigel-coated slide flasks and cultured in differentiation media for 21-28 days. Ectoderm medium: 50% Neurobasal medium, 50% DMEM/F12, 1% N2, 1% B27, 1% Glutamax and 1% Penicillin-Streptomycin; Endoderm medium: Knockout-DMEM, 10% FBS, 1% NEAA, 0.1% β-mercaptoethanol, 1% Glutamax and 1% Penicillin-Streptomycin (all Gibco); Mesoderm medium: Endoderm medium supplemented with 0.5mM ascorbic acid. Cells were analyzed by immunocytochemistry as described above. Antibodies are listed in Supplemental Table M1. Confocal images were taken using Leica TSC SPE/SP5 microscopes.

**Direct differentiation protocols**

**Direct differentiation to mesoderm.** Cells were seeded in a slide-flask with mTeSR medium. Twenty-four hours later (D0), mTeSR was changed by RPMI-B27 minus insulin medium with 8uM Chir. (RPMI-B27 media was composed by RPMI, GLX 1X, P/S 1X, NEEA 1X, B27(-/-) 1X and 2-mercaptoethanol 0,1X). On D1 medium was changed y RPMI-B27 with 2uM Chir. On D3: RPMI-B27 minus insulin with 5uM IWP4. D5: RPMI-B27 without insulin. D7: RPMI-B27 with insulin until D11.

**Direct differentiation to endoderm.** Cells were seeded in matrigel-coated slide flasks with mTeSR medium until 80-90% of confluence and then stablish the protocol as follows:

- D1 - RPMI + P/S + GLX, add 100ng/ml activin A and 0,5uM Chir
- D2 - RPMI + P/S + GLX + 0,2% FBS, add 100ng/ml activi A and bFGF 0,1 µl/ml (100ugr/ml)
- D3 - RPMI + P/S + GLX + 2%FBS activin A + bFGF
- D4 - RPMI + P/S + GLX + B27 add FGF4 1µl/ml (500mM) + 3uM Chir
- D6-D8 – RPMI + P/S + GLX + B27 add FGF4 1µl/ml (500mM) + 3uM Chir

P**rotocol for mix population (neurons and astrocytes)**. EBs were formed and seeded in Matrigel coated slide flasks as described above. The protocol followed:

- 7-10 days with NP-selection medium (Supplemental Table M3)
- 10-15 days NP-Expansion medium (Supplemental Table M3)
- Select and dissect structures neural-like
- Expansion in expansion medium (Supplemental Table M3)
- Seeded in Matrigel coated slide-flasks

Differentiation was analysed by immunocytochemistry as described above. Antibodies are listed in (Supplemental Table M1). Confocal images were taken using Leica TSC SPE/SP5 microscopes.

**Neural Crest Differentiation Media**: DMEM:F12 (Gibco) 1:1; 5mg/mL BSA (Sigma); 500U/mL penicillin/ 500mg/mL streptomycin (Gibco); 2mM GlutaMAX (Gibco); 1x MEM non-essential amino acids (Gibco); 1x trace elements A; 1x trace elements B; 1x trace elements C (Corning); 2-mercaptoethanol (Gibco); 10 µg/mL transferrin (Sigma); 50µg/mL sodium L-ascorbate (Sigma); 10ng/mL heregulin-b1 (PeproTech); 200ng/mL LONG R3 IGFR (PeproTech); 8ng/mL basic fibroblast growth factor 2 (PeproTech), 2µM CHIR9902 (STEMCELL Technologies) and 20µM SB432542 (STEMCELL Technologies).

**Schwann Cell Differentiation Media (SCDM)**: DMEM:F12 (3:1); 500U/ml penicillin/ 500mg/mL streptomycin antibiotics (Gibco); 5µM forskolin (Sigma); 50ng/mL heregulin-b1; 2% N2 supplement (Gibco); 1% FBS (Gibco). Medium was changed twice a week.

**Material and Methods Supplemental Tables**

| **Supplemental Table M1. List of antibodies.** | | | |
| --- | --- | --- | --- |
| **Antibody** | **Supplier** | **Reference** | **Dilution** |
| Rabbit IgG anti-NF2 / Merlin | Abcam | ab109244 | 1:200 |
| Rabbit IgG anti-Vinculin | Abcam | ab129002 | 1:1000 |
| Mouse IgG anti-OCT3/4 | Santa Cruz Biotechnology | Sc-5279 | 1:60 |
| Mouse IgG anti OCT4 (OCT3) | Stem Cell Technologies | #60059 | 1:100 |
| Rabbit IgG anti-SOX2 | Pierce Antibodies | PA1-16968 | 1:100 |
| Goat IgG anti-NANOG | R&D Systems | AF1997 | 1:25 |
| Rat IgM anti-SSEA3 | Hybridoma Bank | MC-631 | 1:3 |
| Mouse IgG anti-SSEA4 | Hybridoma Bank | MC-813-70 | 1:3 |
| Mouse IgM anti TRA-1-81 | Millipore | MAB4381 | 1:400 |
| Goat IgG anti-FOXA2 | R&D Systems | AF2400 | 1:50 |
| Rabbit IgG anti-GATA4 | Santa Cruz Biotechnology | Sc-9053 | 1:50 |
| Mouse IgG anti SMA | Sigma | A5228 | 1:400 |
| Mouse IgM anti-ASA | Sigma | A2172 | 1:400 |
| Rabbit IgG anti GFAP | Dako | Z0334 | 1:500 |
| Mouse IgG anti-TUJ1 | Bio Legend | MMS-435P | 1:500 |
| Mouse IgG anti [NGFR5] to p75 NGF Receptor | Abcam | ab3125 | 1:100 (IF)  1:1000 (FACS) |
| Rabbit IgG anti-S100B | Dako | Z0311 | 1:1000 |
| Mouse IgG anti-AP2 | Thermo Scientific | MA1-872 | 1:50 |
| Rabbit IgG anti-Sox10 | Abcam | ab155279 | 1:50 |
| Mouse IgG anti-HNK1 | SIGMA | C6680 | 1:1000 (FACS) |

| **Supplemental Table M2. List of primers for RT-PCR to detect SeV genome and transgenes set** | | | |
| --- | --- | --- | --- |
| **Target** |  | **Primer** | **Product size (bp)** |
| SeV | *Forward* | GGATCACTAGGTGATATCGAGC | 181 |
|  | *Reverse* | ACCAGACAAGAGTTTAAGAGATATGTATC |  |
| KOS | *Forward* | ATGCACCGCTACGACGTGAGCGC | 528 |
|  | *Reverse* | ACCTTGACAATCCTGATGTGG |  |
| Klf4 | *Forward* | TTCCTGCATGCCAGAGGAGCCC | 410 |
|  | *Reverse* | AATGTATCGAAGGTGCTCAA |  |
| L-Myc | *Forward* | GAGAAGAGGATGGCTACAGAGA | 237 |
|  | *Reverse* | GACGTGCAACTGTGCTATCT |  |

| **Supplemental Table M3. Differentiation media** | | |
| --- | --- | --- |
| **Selection medium (200ml)** | **Expansion medium (200ml)** | **Differentiation medium (200ml)** |
| DMEM-F12 192.8ml | DMEM-F12 191.76ml | DMEM-F12 96.5ml |
| P/S 2ml | P/S 2ml | Neurobasal medium 96.5ml |
| GLX 2ml | GLX 2ml | P/S (1%) 2ml |
| b-mercaptoethanol 200µl | b-mercaptoethanol 200µl | GLX (1%) 2ml |
| NEAA 2ml | NEAA 2 ml | N2 (0.5%) 1ml |
| N2 (0.5%) 1ml | N2 (1%) 2ml | B27 (1%) 2ml |
|  | hFGF 40µl (20ng/ml) |  |

Supplemental Figures

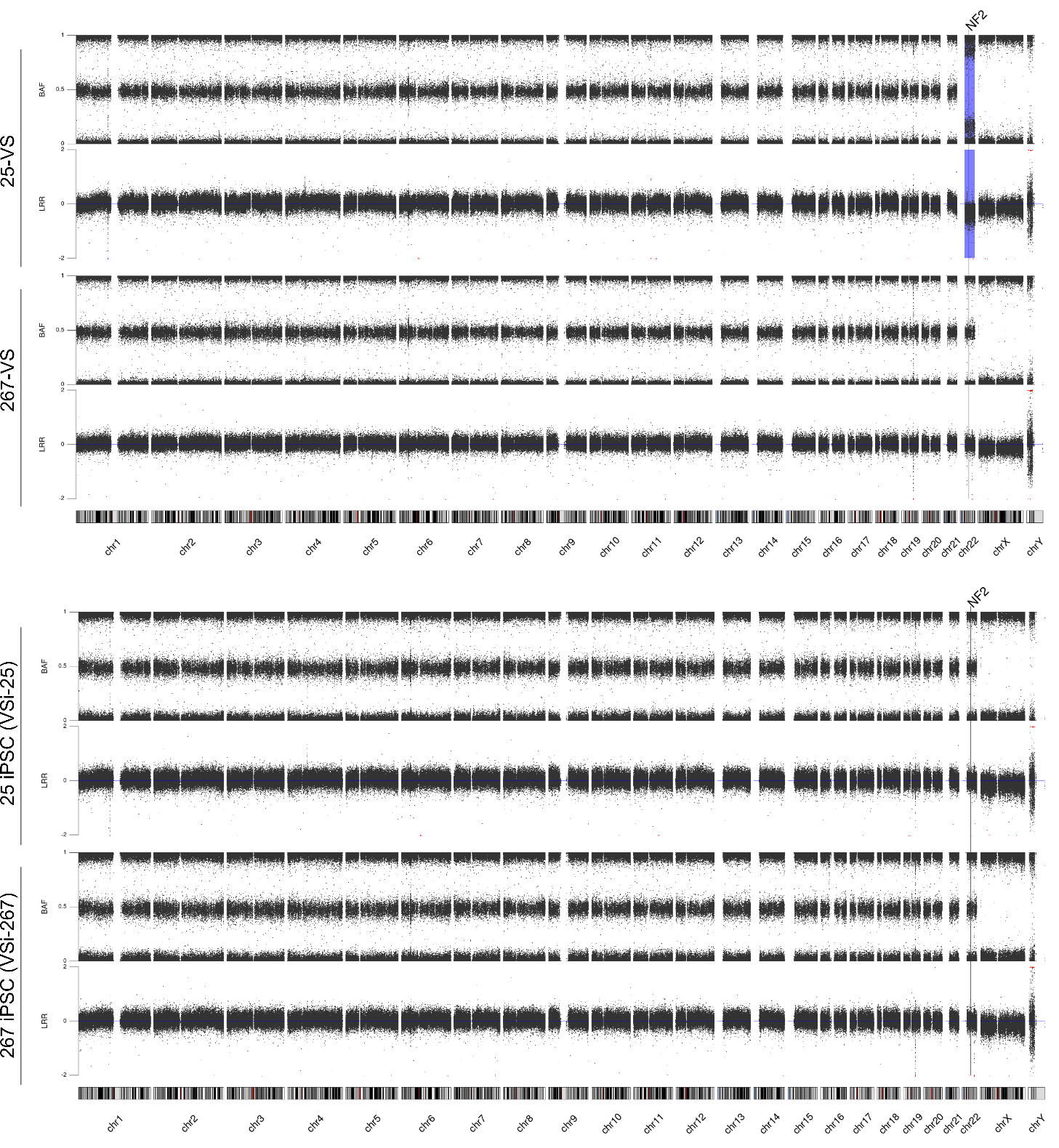
**Supplemental Figure S1 –** SNP-array analysis showed loss of heterozygosity (LOH) on chromosome 22q in VS-25 and no presence of complex rearrangements. VSi-25 and VSi-267 iPSCs SNP-array analysis showed no differences with respect to the tumor of origin, with the exception for VS-25, in which the LOH on chromosome 22q as the second hit was not present in the (+/-) clones obtained from this tumor. BAF: B allele frequency; LRR: Log R Ratio; The blue shaded region denotes the absence of *NF2* due to the LOH.

**Supplemental Figure S2 –** **Characterization of the VSi-245(-/-) iPSC.** (A) Immunochemistry of pluripotency markers NANOG, OCT4, and SOX2 (in green), TRA-1-81, SSEA3 (in red); (B) Morphology of the cell colonies; (C) Immunochemistry to demonstrate the capacity of the iPSCs to in vitro differentiate to the three primary germ layers: mesoderm (ASMA in green and ASA in red), ectoderm (TUJ1 in green and GFAP in red) and endoderm (AFP in green and FOXA2 in red); (D) Alkaline Phosphatase (ALP) Staining; (E) Karyotype at passage 20 (46, XY); (F) Merlin expression analysed by Western Blot. The FiPS line generated from fibroblasts (NF2^+/+^) was used as a control cell line, NPE stands for Normalized Protein Expression. (G) Expression of SeV and transgenes at different passages (1: p8; 2: p40; 3: negative control) assessed by RT-PCR. (H) SNP-array analysis showed no presence of complex rearrangements nor LOH of ch22q.

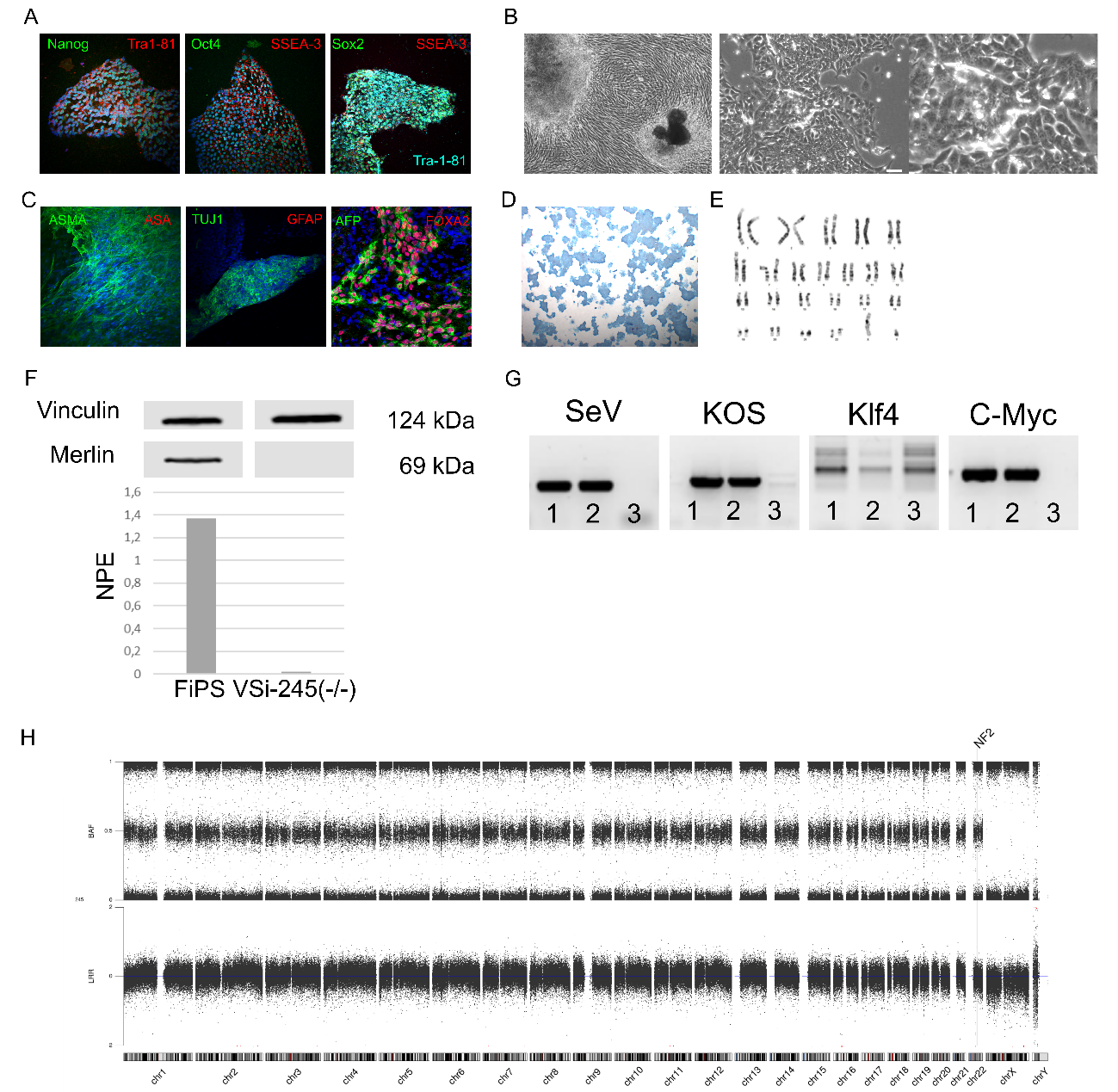

**Supplemental Figure S3 – Characterization of iPSC clones.** (A) Immunochemistry of pluripotency markers NANOG, OCT4, and SOX2 (in green), TRA-1-81, SSEA3 (in red); Scale bar, 75µM; Cell nuclei were stained with DAPI; (B) Alkaline Phosphatase (ALP) Staining; (C) Karyotype of lines at passage 20 (46, XY).

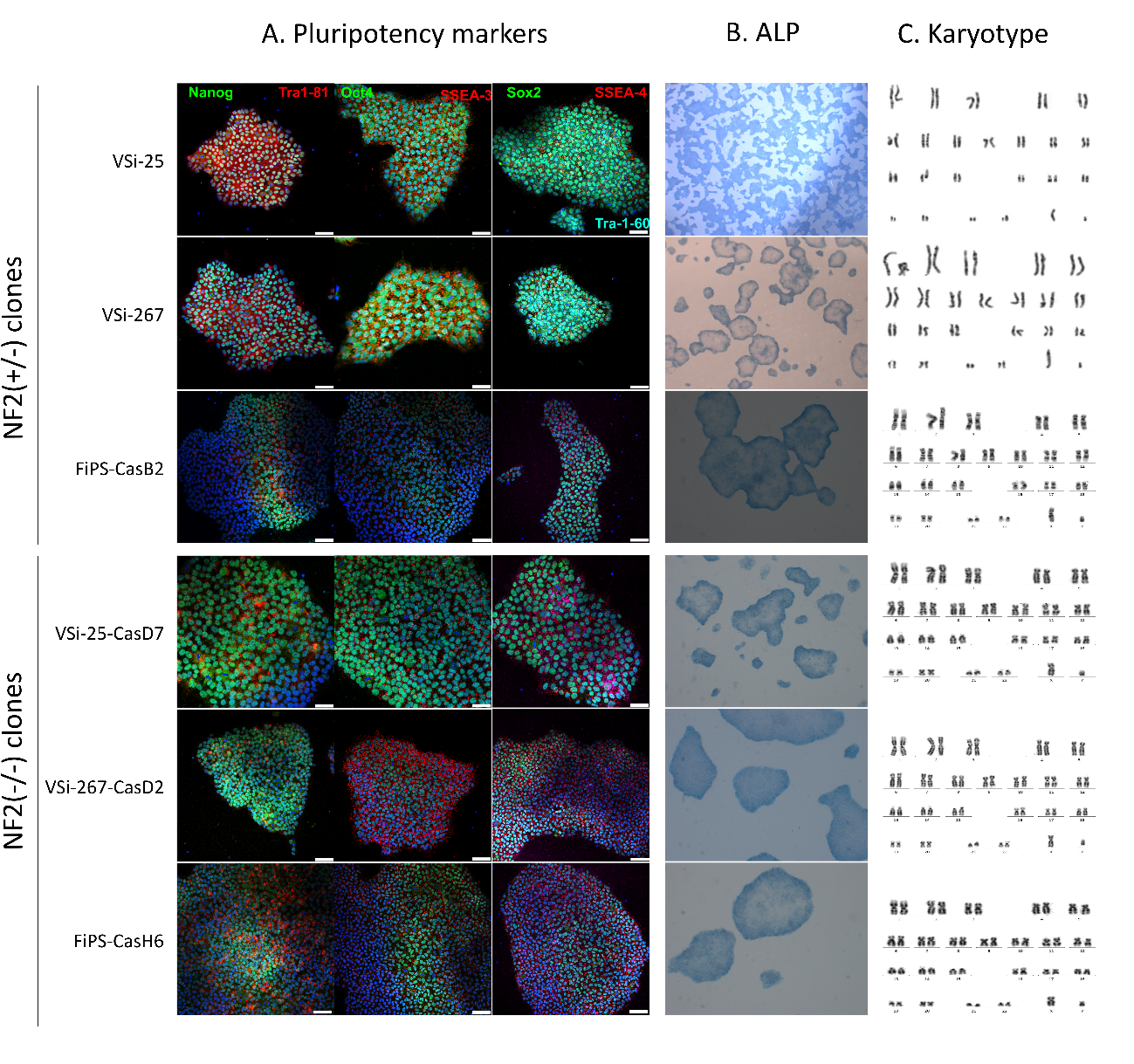

**Supplemental Figure S4** –**Phase-contrast images of *NF2* (+/+), *NF2* (+/-) and *NF2* (-/-) cell lines during SC differentiation**. *NF2* deficient cells did not show capacity to maintain attachment to the cell culture already after 5 days under the SC differentiation media. Scale bar, 75µM.

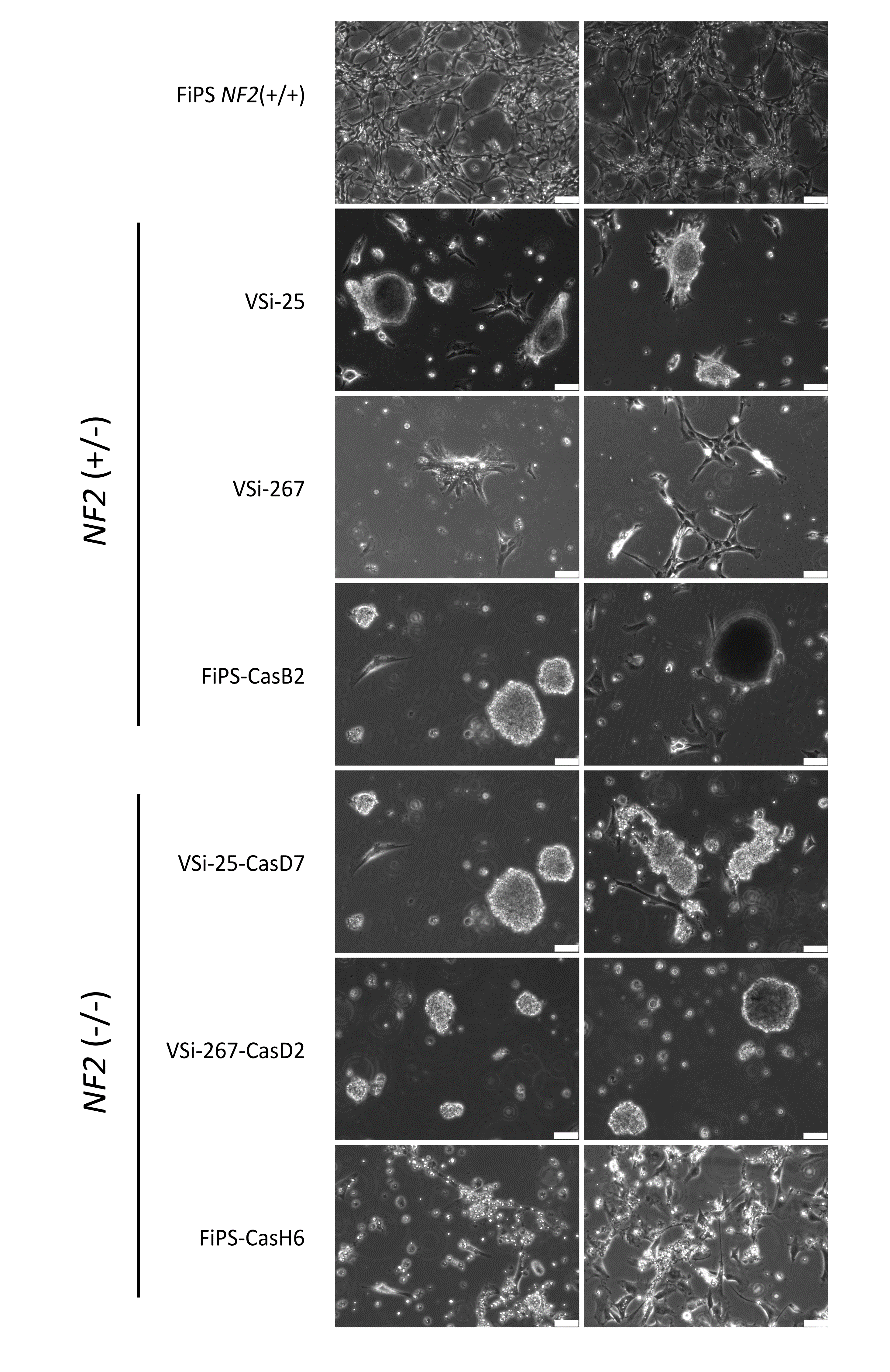

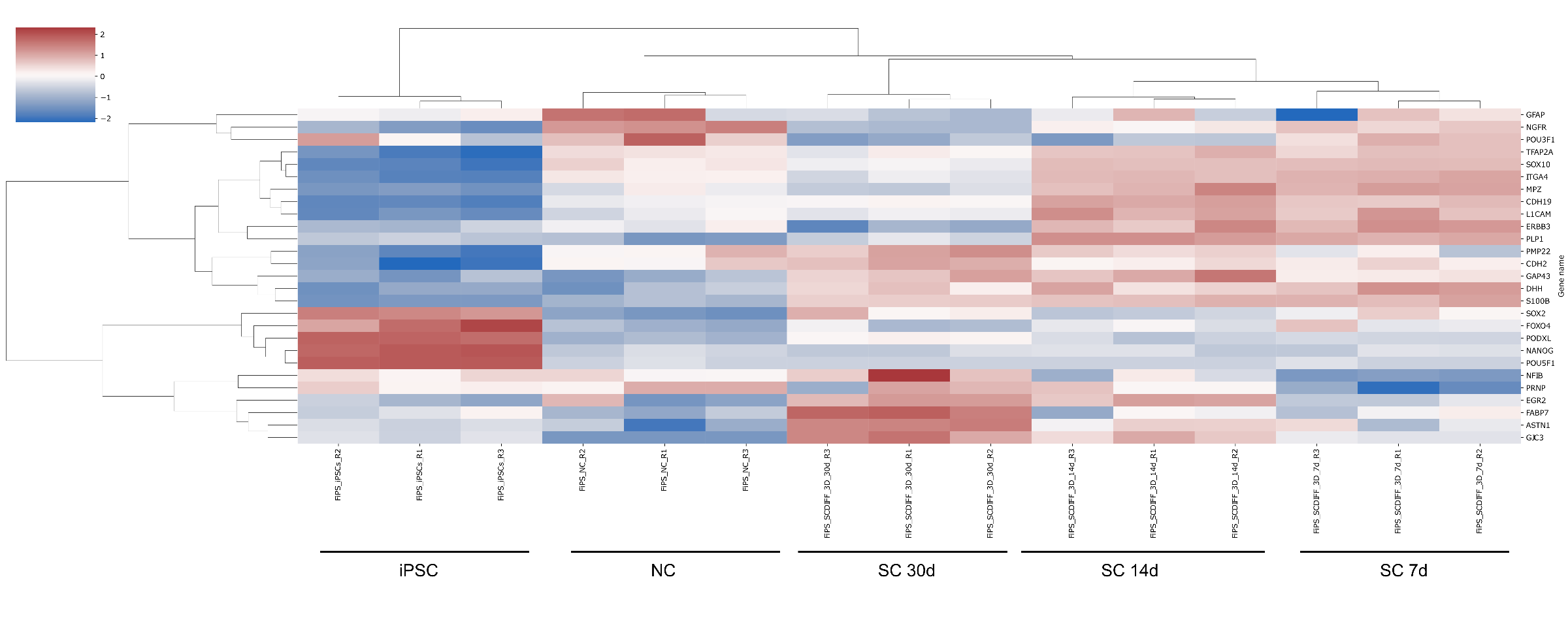
**Supplemental Figure S5**. Heatmap of the control *NF2*(+/+) line (FiPS) in the 3D differentiation protocol, representing expression of genes expressed over the stages of the differentiation protocol. Data shown is from three independent differentiation experiments. 7d, 14d and 30d stand for 7, 14 and 30 days under SC differentiation conditions, respectively.

**Supplemental Figure S6** – **Phase contrast images of *NF2*(+/-) and *NF2*(-/-) 3D cultures at day 7 of SC differentiation in 3D cultures.** No spheroids could be generated from the FiPS-CasB2 (*NF2* +/-) NC derived cells, Scale bar, 75µM (left panel) and 250µM (right panel).

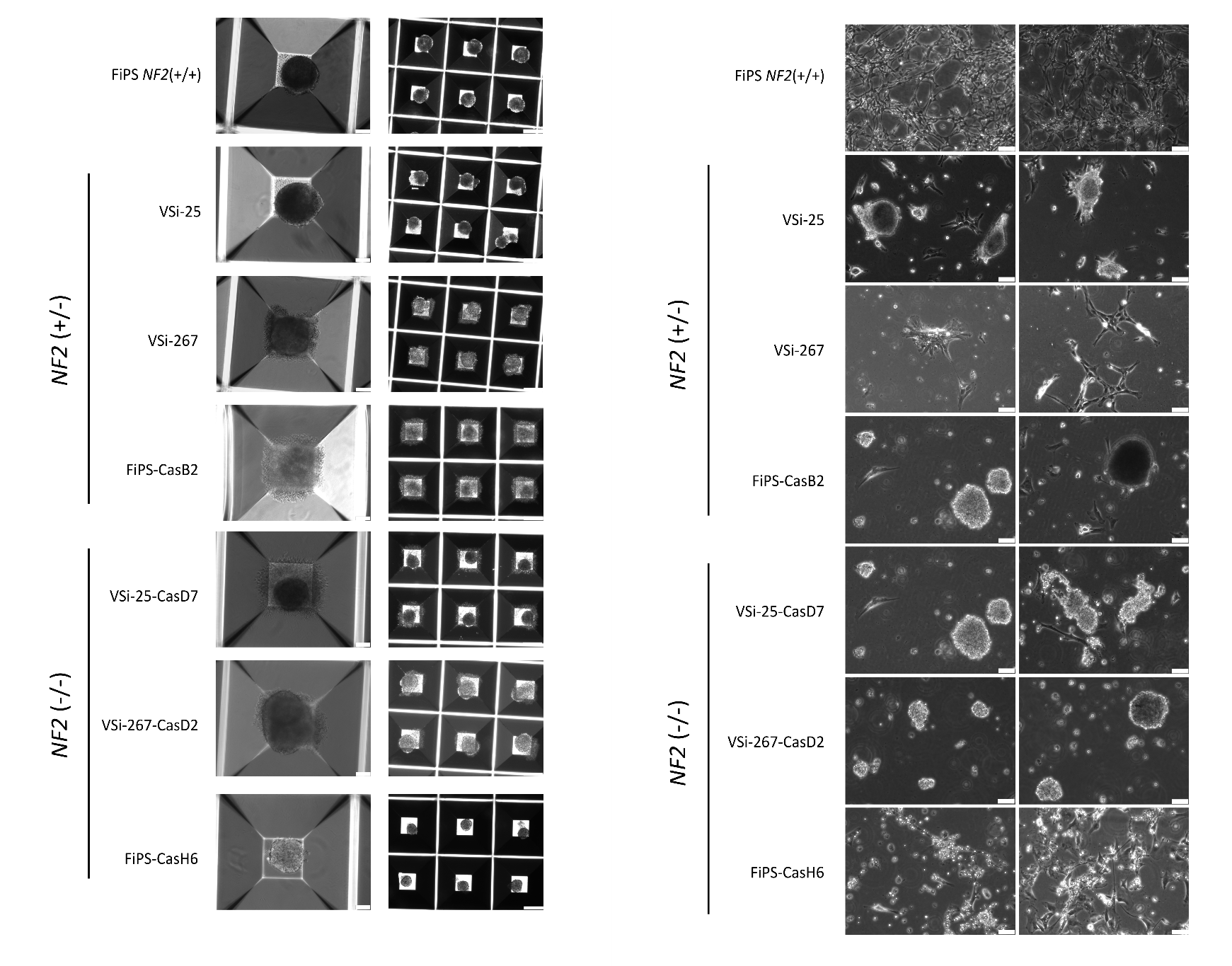

**Supplemental Figure S7**. *In vitro* NC-SC expression roadmap of the control line (FiPS, *NF2*(+/+)) in the 2D differentiation protocol and the *NF2*(-/-) lines in the 3D differentiation protocol. Data shown is from three independent differentiation experiments. 7d and 14d stand for 7 and 14 days under SC differentiation conditions, respectively.
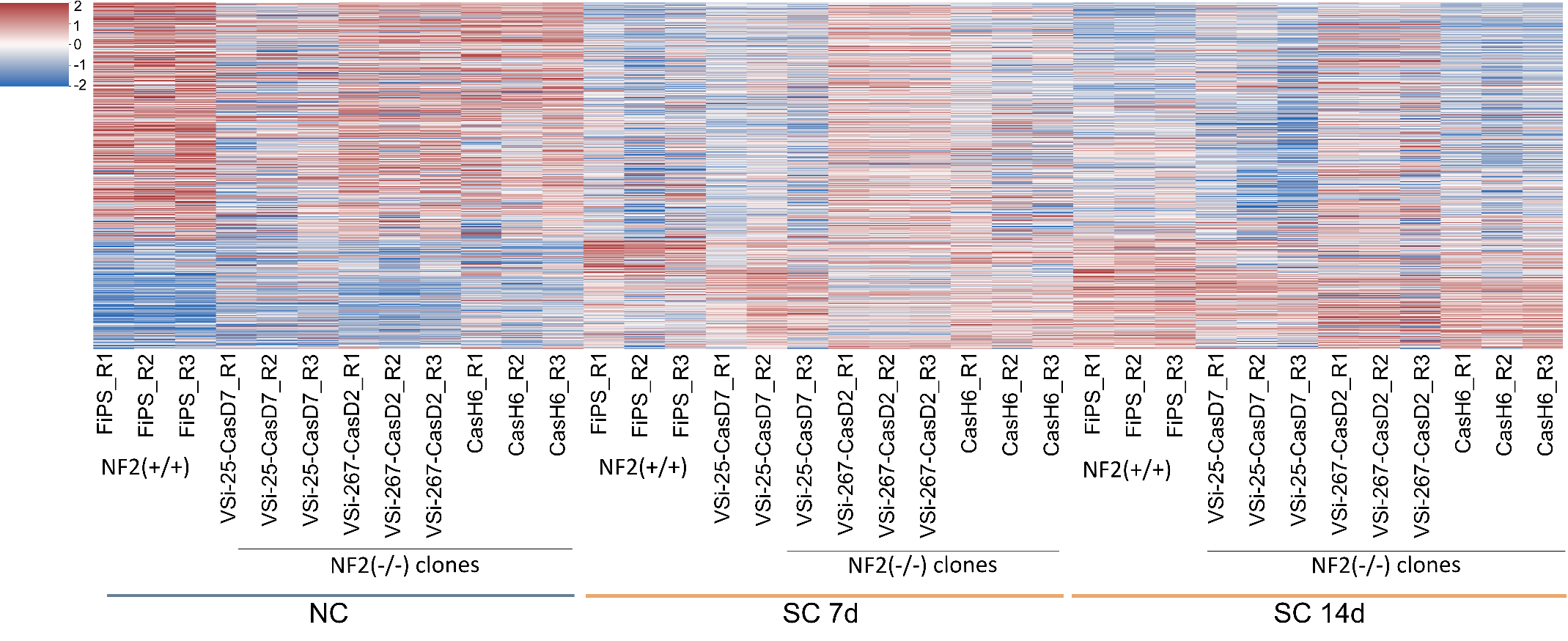

**Supplemental Figure S8.** (A) Volcano plot for DE genes between *NF2*(+/-) vs *NF2*(+/+). X-axis shows Log2-fold changes between conditions. Y-axis represents the p-value in –log10 scale. padj < 0.01 logfc =1 (B) GSEA analysis between *NF2*(+/-) vs *NF2*(+/+) are shown. Only Hallmark pathways with FDR<0.05 are displayed. Green bars on X-axis account for the enrichment score on each of them. (C) Volcano plot for DE genes between *NF2*(-/-) vs *NF2*(+/+). (D) GSEA analysis between *NF2*(-/-) vs *NF2*(+/+).

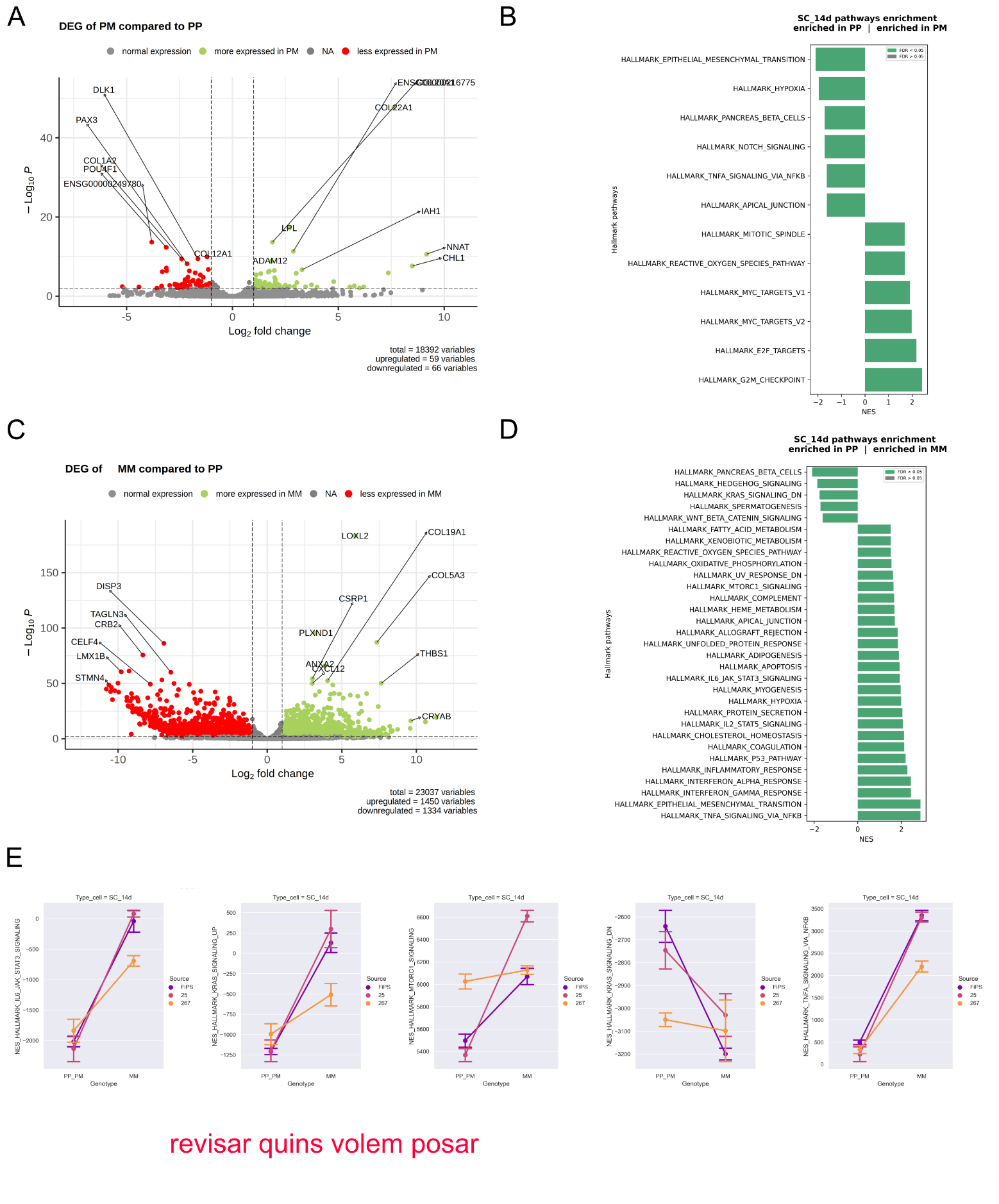

**Supplemental Figure S9**. (A) T-test was performed for each individual comparison between genotypes. Bars express mean normalized expression ±SD from three independent experiments. Significant comparisons are shown as *<0.05; **<0.01, ***<0.001. (B) PCA analysis over the 2000 genes with highest SD. Components 1 and 2 are shown in left panel, and components 2 and 3 in the right one. Explained variance is indicated for each of them. (C) EnrichR significant pathways for schwannoma and *NF2*(+/+) genes upregulated in both groups. X-axis correspond to –log10 p-values for each of them.

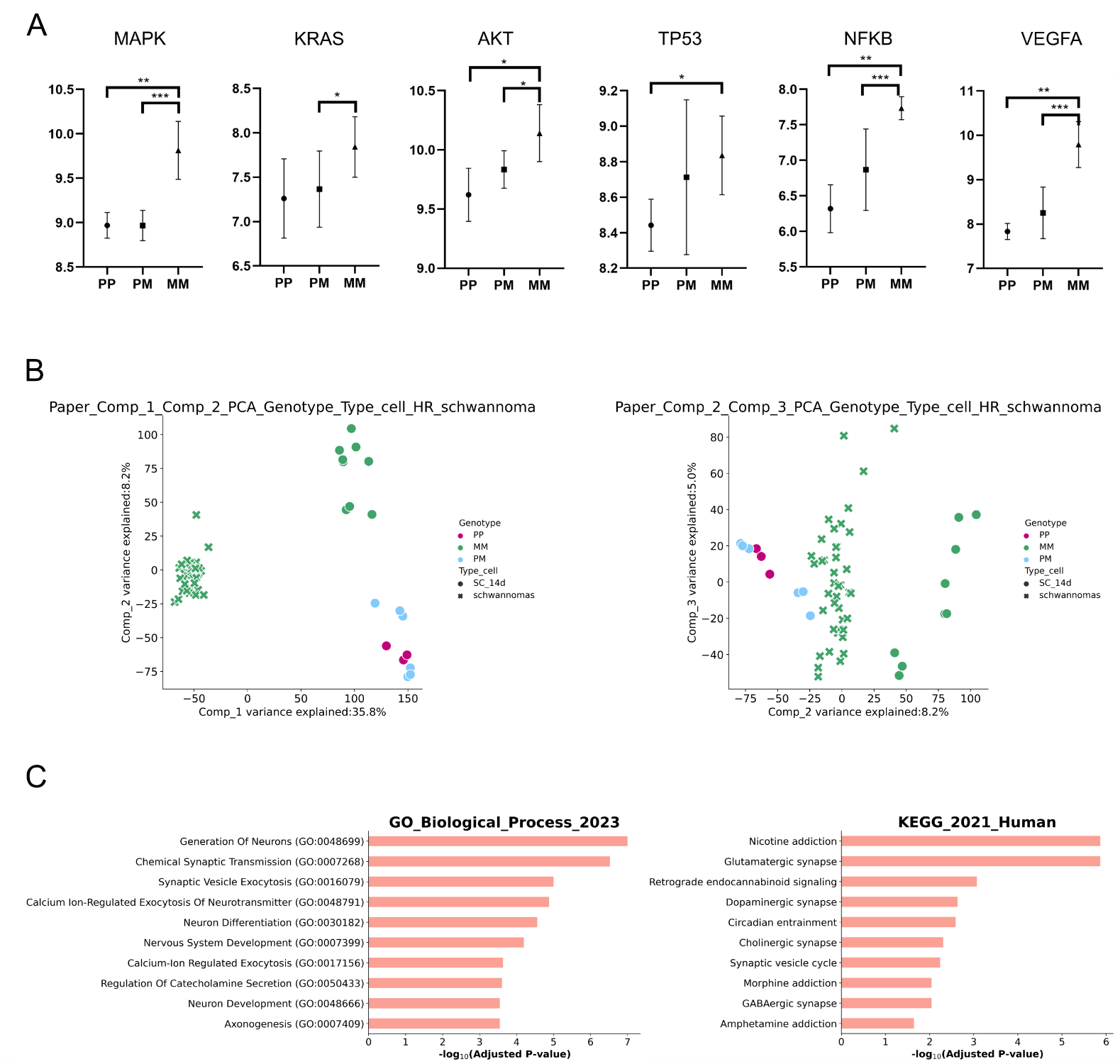

**Supplemental Tables**

| Supplemental Table S1. | | | |
| --- | --- | --- | --- |
| Patient ID | 25 | 245 | 267 |
| Sex | XY | XY | XY |
| Diagnostic | NF2 | Mosaic NF2 | NF2 |
| Age at diagnosis | 47 | 23 | 14 |
| Tumor load | BVS | Unilateral VS | BVS, Peripheral SC and multiple intraspinal SC |
| Number of interventions | 2 | 1 | 3 |
| Age at first intervention | 47 | 23 | 14 |
| **VS ID** | VS-25 | VS-245 | VS-267 |
| ***NF2* Germline Mutation** | g.83045A>G | g.73341insG | g.62758C>T |
| ***NF2* Somatic Mutation** | LOH | g.76297C>T | g.73394G>A |

| **Supplemental Table S2. VSs Reprogramming information** | | | | | | |
| --- | --- | --- | --- | --- | --- | --- |
|  | | | | **iPSCs clones genotype** | | |
| **Patient ID** | **VS ID** | **Method** | **Number of Clones Analysed** | ***NF2*(+/+)** | ***NF2*(+/-)** | ***NF2*(-/-)** |
| 25 | VS-25 | SeV | 38 | 0 | 38 | 0 |
| 267 | VS-267 | SeV | 12 | 0 | 12 | 0 |
| 245 | VS-245 | SeV | 10 | 1 | 0 | 9 |

| **Supplemental Table S3. *In vitro* iPSC differentiation assays** | | | | | | |
| --- | --- | --- | --- | --- | --- | --- |
| ***NF2* Genotype** | **+/-** | **+/-** | **+/-** | **-/-** | **-/-** | **-/-** |
|  |  |  | **NF2-FiPS-PM-B2** | **NF2-FiPS-MM-H6** | **NF2-25iPS-MM-D7** | **NF2-267iPS-MM-D2** |
| ***In vitro* differentiation (EBs) Ectoderm** | Tuj1 +  GFAP + | Tuj1 +  GFAP + | Tuj1 +  GFAP + | Tuj1 +  GFAP + | EBs do not succeed | EBs do not succeed |
| ***In vitro* differentiation (EBs) Endoderm** | AFP +  FOXA2 + | AFP +  FOXA2 + | AFP +  FOXA2 + | EBs do not succeed | EBs do not succeed | EBs do not succeed |
| ***In vitro* differentiation (EBs) Mesoderm** | ASMA +  GATA4 + | ASMA +  GATA4 + | ASMA +  GATA4 + | EBs do not succeed | EBs do not succeed | EBs do not succeed |
| **Direct differentiation Ectoderm** |  |  | **-** |  | Tuj1 + GFAP – PAX6 + \| | Tuj1 +  GFAP –  PAX6 + |
| **Direct differentiation Endoderm** |  |  | **-** | AFP +  FOXA2 +  SOX17 + | AFP +  FOXA2 +  SOX17 + | AFP +  FOXA2 +  SOX17 + |
| **Direct differentiation Mesoderm** |  |  | **-** | ASMA +  GATA4 + | ASMA +  GATA4 + | ASMA +  GATA4 + |

| **Supplemental Table S4. WES analysis of the CRISPR-generated lines** | | | | | | |
| --- | --- | --- | --- | --- | --- | --- |
| **FiPS-CasB2** | | | | | | |
| **Gene_Symbol** | **Variant_Classification** | **Variant_Type** | **Genome_Change** | **cDNA_Change** | **Protein_Change** | **SwissProt_acc_Id** |
| DNAJC10 | Missense_Mutation | SNP | g.chr2:183582950C>A | c.137C>A | p.A46E | Q8IXB1 |
| LIMD1 | Missense_Mutation | SNP | g.chr3:45637329G>A | c.958G>A | p.G320S | Q9UGP4 |
| HMGXB3 | Intron | SNP | g.chr5:149425299T>A | c.e16+51T>A |  | Q12766 |
| GBA2 | Missense_Mutation | SNP | g.chr9:35748662G>A | c.40C>T | p.P14S | Q9HCG7 |
| B3GNTL1 | Intron | DEL | g.chr17:80993071_80993073delTCG | c.e10+95CGAA>A |  | Q67FW5 |
| KANK3 | Missense_Mutation | SNP | g.chr19:8400524G>A | c.187C>T | p.R63C | Q6NY19 |
| NF2 | Frame_Shift_Del | DEL | g.chr22:30032834_30032835delAC | c.209_210delAC | p.T71fs | P35240 |
| **FiPS-CasH6** | | | | | | |
| **Gene_Symbol** | **Variant_Classification** | **Variant_Type** | **Genome_Change** | **cDNA_Change** | **Protein_Change** | **SwissProt_acc_Id** |
| YRDC | Missense_Mutation | SNP | g.chr1:38272621C>T | c.532G>A | p.A178T | Q86U90 |
| EIF2B3 | Intron | SNP | g.chr1:45443916T>C | c.e10-71A>G |  | Q9NR50 |
| CRNN | Intron | SNP | g.chr1:152384514C>A | c.e2-58G>T |  | Q9UBG3 |
| OR10J1 | Missense_Mutation | SNP | g.chr1:159409651C>A | c.70C>A | p.Q24K | P30954 |
| BLOC1S4 | Silent | SNP | g.chr4:6718341C>A | c.405C>A | p.I135I | Q9NUP1 |
| THAP6 | 5'UTR | SNP | g.chr4:76441986C>A |  |  | Q8TBB0 |
| CHD1 | Frame_Shift_Del | DEL | g.chr5:98236745delT | c.629delA | p.K210fs | O14646 |
| MTCH1 | Intron | SNP | g.chr6:36944179G>T | c.e6+686C>A |  | Q9NZJ7 |
| ANKRD30A | Missense_Mutation | SNP | g.chr10:37490239G>A | c.3212G>A | p.S1071N | Q9BXX3 |
| AGAP2 | Missense_Mutation | SNP | g.chr12:58125611C>T | c.926G>A | p.R309H | Q99490 |
| TPP2 | Missense_Mutation | SNP | g.chr13:103328667C>T | c.3562C>T | p.H1188Y | P29144 |
| IQGAP1 | Intron | SNP | g.chr15:90986819G>A | c.e9+109G>A |  | P46940 |
| ATCAY | Missense_Mutation | SNP | g.chr19:3909578G>A | c.742G>A | p.G248S | Q86WG3 |
| EVI5L | Frame_Shift_Del | DEL | g.chr19:7918216delC | c.1143delC | p.R383fs | Q96CN4 |
| NF2 | Frame_Shift_Del | DEL | g.chr22:30032834_30032835delAC | c.209_210delAC | p.T71fs | P35240 |
| NF2 | Frame_Shift_Ins | INS | g.chr22:30032835_30032836insA | c.210_211insA | p.T71fs | P35240 |
| **VSi-25-CasD7** | | | | | | |
| **Gene_Symbol** | **Variant_Classification** | **Variant_Type** | **Genome_Change** | **cDNA_Change** | **Protein_Change** | **SwissProt_acc_Id** |
| WDR75 | Intron | SNP | g.chr2:190334990A>G | c.e17+19A>G |  | Q8IWA0 |
| TSEN2 | Missense_Mutation | SNP | g.chr3:12571316C>T | c.1192C>T | p.P398S | Q8NCE0 |
| IGSF10 | Missense_Mutation | SNP | g.chr3:151166779C>T | c.990G>A | p.M330I | Q6WRI0 |
| AMTN | Intron | SNP | g.chr4:71384605C>A | c.e2+57C>A |  | Q6UX39 |
| PAPSS1 | Intron | SNP | g.chr4:108565897T>C | c.e3-61A>G |  | O43252 |
| TRGV1 | RNA | SNP | g.chr7:38407419G>C | c.119C>G |  |  |
| MAGI2 | Intron | DEL | g.chr7:77702356delG | c.e2-5909CA>A |  | Q86UL8 |
| MAGI2 | Intron | SNP | g.chr7:77702357C>A | c.e2-5907G>T |  | Q86UL8 |
| CLCN1 | Splice_Site | SNP | g.chr7:143028744C>T | c.1165C>T | p.H389Y | P35523 |
| DLC1 | Intron | SNP | g.chr8:13133754C>A | c.e14-29024G>T |  | Q96QB1 |
| TMEM126B | Missense_Mutation | SNP | g.chr11:85345187A>G | c.261A>G | p.I87M | Q8IUX1 |
| APPL2 | Intron | SNP | g.chr12:105623044G>T | c.e20+43C>A |  | Q06481 |
| YY1 | Nonsense_Mutation | SNP | g.chr14:100706206C>T | c.625C>T | p.Q209* | P25490 |
| SLC24A1 | Missense_Mutation | SNP | g.chr15:65938015G>T | c.2206G>T | p.D736Y | O60721 |
| TICRR | Intron | SNP | g.chr15:90128887C>A | c.e4-52C>A |  | Q7Z2Z1 |
| LGALS13 | Intron | SNP | g.chr19:40095199C>A | c.e2-43C>A |  | Q9UHV8 |
| MARK4 | Intron | SNP | g.chr19:45767930C>G | c.e5-12C>G |  | Q96L34 |
| CST2 | Missense_Mutation | SNP | g.chr20:23807073C>A | c.225G>T | p.E75D | P09228 |
| NF2 | Frame_Shift_Del | DEL | g.chr22:30032834_30032835delAC | c.209_210delAC | p.T71fs | P35240 |
| NF2 | Frame_Shift_Ins | INS | g.chr22:30032835_30032836insA | c.210_211insA | p.T71fs | P35240 |
| **VSi-267-CasD2** | | | | | | |
| **Gene_Symbol** | **Variant_Classification** | **Variant_Type** | **Genome_Change** | **cDNA_Change** | **Protein_Change** | **SwissProt_acc_Id** |
| TTC34 | Silent | SNP | g.chr1:2706547C>A | c.1149G>T | p.L383L | A8MYJ7 |
| SIDT1 | Intron | SNP | g.chr3:113342242C>T | c.e22-32C>T |  | Q9NXL6 |
| TRIM4 | Intron | SNP | g.chr7:99490380A>T | c.e1+11T>A |  | Q9C037 |
| TRBV24OR9-2 | RNA | SNP | g.chr9:33649340C>A | c.e2+29C>A |  |  |
| DPH1 | Intron | SNP | g.chr17:1943961G>A | c.e9+62G>A |  | Q9BZG8 |
| CCDC178 | Intron | SNP | g.chr18:30672889A>T | c.e3+15T>A |  | Q5BJE1 |
| NF2 | Frame_Shift_Del | DEL | g.chr22:30032834_30032835delAC | c.209_210delAC | p.T71fs | P35240 |
